# Biophysical characterization of the human K0513 protein

**DOI:** 10.1101/2020.06.18.158949

**Authors:** Ndivhuwo Nemukondeni, Afolake Arowolo, Addmore Shonhai, Tawanda Zininga, Adélle Burger

**Affiliations:** Department of Biochemistry, School of Mathematical & Natural Sciences, University of Venda, Private Bag X5050, Thohoyandou, 0950, South Africa; Department of medicine, University of Cape Town, Observatory,7925, South Africa; Department of Biochemistry, Stellenbosch University, Stellenbosch, 7602, South Africa

**Keywords:** K0513, recombinant protein, SBF2/MTMR13, GTP binding, GEF, Rab3a

## Abstract

Glioblastoma multiforme (GBM) is an aggressive grade IV primary malignant tumour which accounts for 78 % of all brain tumours. *K0513* is a GBM biomarker that is upregulated in the invasive phenotype. *K0513* is expressed ubiquitously and is reportedly enriched in the cerebral cortex of the brain. *K0513* is further implicated in signalling pathways involving neuroplasticity, cytoskeletal regulation and apoptosis. The protein encoded by *K0513* is a prospective biomarker for pancreatic cancer prognosis. However, the gene product of *K0153* is not well characterised. This study focused on structure-function characterisation of human K0513 protein. To this end, we employed bioinformatics analysis and biophysical approaches to characterize the protein. *In silico* structural characterisation of the human K0513 protein suggests the presence of a SET binding factor 2 (SBF2) domain and a transmembrane region. The SBF2 domain is found in the Myotubularin-related protein 13 (MTMR13), which may function as a nucleotide exchange factor for the RAS-associated GTPase, Rab28. K0513 was predicted to interact with RAS-associated GTPase, Rab3a. Secondary structure prediction revealed K0513 to be predominantly α-helical in nature. The predicted three-dimensional model of K0513 showed a globular fold, suggesting that the protein is water-soluble. K0513 was heterologously expressed in *E. coli* XL1-Blue cells and subsequent purification yielded 80 % soluble protein. Biophysical characterisation using tryptophan-based fluorescence spectroscopy and limited proteolysis showed the conformation of K0513 is mostly unperturbed in the presence of nucleotides. Interestingly, K0513 was detected in lung carcinoma, fibrosarcoma and cervical adenocarcinoma cells, supporting its possible role in cancer signalling pathways.

## 1. INTRODUCTION

Glioblastoma multiforme (GBM) is one of the most lethal adult brain tumours. GBM is a fast-growing and nonmetastatic cancer that may invade tissues of the brain and spine. GBM patients have a very poor prognosis with a median survival of ∼10 months [1], a 5,1 % survival rate over five years, and a 37,2 % survival rate over a single year [2]. GBM cells are highly complex and heterogeneous, and display diverse gene expression profiles. Various diagnostic and therapeutic glioblastoma biomarkers are identified by gene expression profiling. Extensive alterations of certain genes amongst the various types of glioblastomas are known to disrupt key signalling pathways [3, 4]. One such gene is the novel *K0513. K0513* was reported to be upregulated in an invasive form of GBM (iGBM) [5]. *K0513* is ubiquitously expressed and enriched in the cerebral region of the brain [6], and an upregulation of this gene is correlated with poor disease prognosis in iGBM and pancreatic cancer [5,7]. In addition, an altered expression of *K0513*, due to atypical methylation patterns, is associated with non-survival of neuroblastoma patients [8]. The Human Protein Atlas (Pathology Atlas) reports that the K0513 protein is expressed in 20 different cancers [9]. Although K0513 is not well characterized, it is suggested that K0513 is associated with tumour progression [10]. K0513 is also reported to interact with proteins involved in signalling pathways of neuroplasticity, apoptosis and cytoskeletal regulation, such as kidney and brain expressed protein (KIBRA) or WW domain-containing protein 1 (WWC1); Hematopoietic Cell-Specific Lyn Substrate 1 (HCLS1)-associated protein X-1 (HAX-1), and integrator complex subunit 4 (INTS4), respectively [6]. K0513 seems to be associated with aggressive cancer forms including iGBM, pancreatic cancer and neuroblastoma, and because of this, its interactome would be important as an indicator of its functional pathway in cancer progression. Thus, there is a need to characterize K0513. Based on structure-function features, the association of K0513with aggressive cancers could serve as a direct or indirect biomarker of iGBM and other invasive cancers and hence, aid in the design or repurposing of novel or existing drugs respectively. The paucity of information on the structure-function features of this apparently important molecule makes its characterization a priority.

## 2. MATERIALS AND METHODS

### 2.1. Materials

Unless otherwise stated, the materials used in this study were procured from Merck KGaA and Sigma-Aldrich (Darmstadt, Germany), Melford Laboratories (Suffolk, UK), and VWR International (Pennsylvania, USA). Mouse raised monoclonal anti-polyhistidine (His) antibodies, and goat anti-mouse IgG horseradish peroxidase (HRP) conjugated antibodies were purchased from ThermoFisher Scientific (Massachusetts, USA). Rabbit raised polyclonal anti-K0513 antibodies, and goat anti-rabbit IgG HRP conjugated antibodies were purchased from Abcam (Cambridge, UK; AB121430).

### 2.2. Bioinformatics analysis of human K0513

Amino acid sequences of K0513 (K0513a: NCBI accession number: NP_001273494.1), and isoforms (K0513b: NCBI accession number:NP_001273495.1; K0513c: NCBI accession number: PN_001273695.1) and homologues from *Pan troglodytes* (PtK0513, NCBI accession number: XP_003952958.1), *Xenopus tropicalis* (XtK0513, NCBI accession number: NP_001096454.1), *Canis lupus* (ClK0513, NCBI accession number: XP_005620695.1), *Bos taurus* (BtK0513, NCBI accession number: XP_005218484.1), *Mus musculus* (MmK0513, NCBI accession number: NP_001157232.1), *Rattus norvegicus* (RnK0513, NCBI accession number: NP_001100906.1), *Gallus gallus* (GgK0513, NCBI accession number: XP_001235240.2), *Danio rerio* (DrK0513, NCBI accession number: NP_001002193.1), and *Macaca mulatta* (MamK0513, NCBI accession number: XP_001113080.1), and 40 orthologues (NCBI accession numbers shown in Fig 1) were retrieved from the National Centre for Biotechnology Information [11]. Phylogenetic tree analysis was used to establish orthology and a multiple sequence alignment was generated using the COBALT Multiple Alignment Tool and Neighbour Joining analysis was performed. The rooted tree was generated using a maximum sequence difference of 0,7, and a Kimura protein distance model. To the best of our knowledge, none of the homologues nor orthologues have been characterized. The multiple sequence alignment of K0513 and homologues was generated using Clustal Omega [12] and shaded using Boxshade. K0513 protein secondary structure was predicted by generating a multiple sequence alignment using PROMALS3D [13]. The analysis was confirmed using PHYRE^2^ [14], which predicts the percentage composition of alpha-helices, beta-sheets, and loops. Furthermore, the hydrophobicity and amphipathicity of K0513 was predicted using Biotools Web-based Hydropathy, Amphipathicity and Topology (WHAT) [15] to predict whether the protein makes contact with a membrane. The amino acid composition of K0513 was computed using PEPSTAT module integrated in the EMBOSS software [16]. Physicochemical prediction of the protein was conducted using the Expasy’s ProtParam server [17]. A three-dimensional structure of K0513 was generated using PHYRE^2^ and the resulting PDB file was visualized using Biovia Discovery studio [18]. An online Integrated Interaction Database (IID) was launched [19] using the K0513 sequence as identifier (UNIPROT ID; O60268) to generate the predicted interactome.

**Fig 1.**
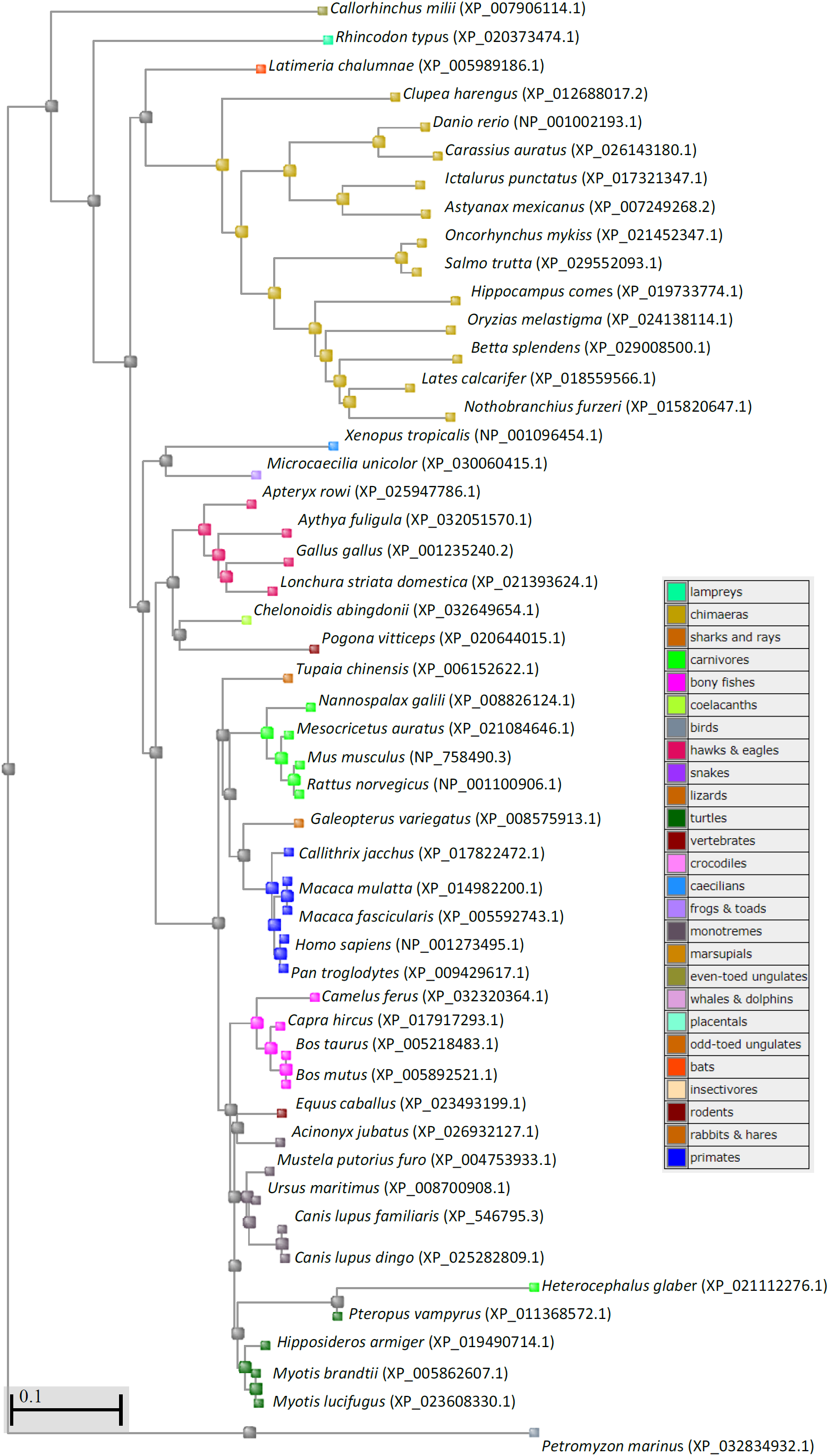
Phylogenetic analysis of homologues and orthologues of human K0513. K0513 amino acid accession numbers were obtained from NCBI. *Callorhinchus milii* K0513 (NCBI accession number: XP_007906114.1); *Rhincodon typus* K0513 (NCBI accession number: XP_020373474.1); *Latimeria chalumnae* (NCBI accession number: XP_005989186.1); *Clupea harengus* (NCBI accession number: XP_012688017.2); *Danio rerio* (NCBI accession number: NP_001002193.1); *Carassius auratus* (NCBI accession number: XP_026143180.1); *Ictalurus punctatus* (NCBI accession number: XP_017321347.1); *Astyanax mexicanus* (NCBI accession number: XP_007249268.2); *Oncorhynchus mykiss* (NCBI accession number: XP_021452347.1); *Salmo trutta* (NCBI accession number: XP_029552093.1); *Hippocampus comes* (NCBI accession number: XP_019733774.1); *Oryzias melastigma* (NCBI accession number: XP_024138114.1); *Betta splendens* (NCBI accession number: XP_029008500.1); *Lates calcarifer* (NCBI accession number: XP_018559566.1); *Nothobranchius furzeri* (NCBI accession number: XP_015820647.1); *Xenopus tropicalis* (NCBI accession number: NP_001096454.1); *Microcaecilia unicolor* (NCBI accession number: XP_030060415.1); *Apteryx rowi* (NCBI accession number: XP_025947786.1); *Aythya fuligula* (NCBI accession number: XP_032051570.1); *Gallus gallus* (NCBI accession number: XP_001235240.2); *Lonchura striata domestica* (NCBI accession number: XP_021393624.1); *Chelonoidis abingdonii* (NCBI accession number: XP_032649654.1); *Pogona vitticeps* (NCBI accession number: XP_020644015.1); *Tupaia chinensis* (NCBI accession number: XP_006152622.1); *Nannospalax galili* (NCBI accession number: XP_008826124.1); *Mesocricetus auratus* (NCBI accession number: XP_021084646.1); *Mus musculus* (NCBI accession number: NP_758490.3); *Rattus norve*gicus (NCBI accession number: NP_001100906.1); *Galeopterus variegatus* (NCBI accession number: XP_008575913.1); *Callithrix jacchus* (NCBI accession number: XP_017822472.1); *Macaca mulatta* (NCBI accession number: XP_014982200.1); *Macaca fascicularis* (NCBI accession number: XP_005592743.1); *Homo sapiens* (NCBI accession number: NP_001273495.1); *Pan troglodytes* (NCBI accession number: XP_009429617.1); *Camelus ferus* (NCBI accession number: XP_032320364.1); *Capra hircus* (NCBI accession number: XP_017917293.1); *Bos taurus* (NCBI accession number: XP_005218483.1); *Bos mutus* (NCBI accession number: XP_005892521.1); *Equus caballus* (NCBI accession number: XP_023493199.1); *Acinonyx jubatus* (NCBI accession number: XP_026932127.1); *Mustela putorius furo* (NCBI accession number: XP_004753933.1); *Ursus maritimus* (NCBI accession number: XP_008700908.1); *Canis lupus familiaris* (NCBI accession number: XP_546795.30); *Canis lupus dingo* (NCBI accession number: XP_025282809.1); *Heterocephalus glaber* (NCBI accession number: XP_021112276.1); *Pteropus vampyrus* (NCBI accession number: XP_011368572.1); *Hipposideros armiger* (NCBI accession number: XP_019490714.1*); Myotis brandtii* (NCBI accession number: XP_005862607.1); *Myotis lucifugus* (NCBI accession number: XP_023608330.1). A multiple sequence alignment was generated using the COBALT multiple alignment tool and Neighbour Joining analysis was performed using COBOLT RID. The rooted tree was generated using a maximum sequence difference of 0,7, and a Kimura protein distance model.

### 2.3. Construction of human K0513 expression vector

The complementary DNA (cDNA) sequence coding for the human K0513 isoform a protein was codon harmonised using the OptimumGene™algorithm for bacterial system expression. This cDNA was subsequently amplified by PCR using primers that introduced *Nco*I and *Bgl*II restriction sites at 5’ and 3’ end of the amplified gene respectively. The 1241 bp DNA fragment comprising *K0513* was subcloned into the expression vector pQE60 between the *Nco*I and *Bgl*II restriction sites to generate a C-terminally 6xHis tagged protein. The pQE60-K0513 construct was synthesized by GenScript (Piscataway, USA). The C-terminal His-tagged pQE60-K0513 construct was verified by sequencing and restriction digestion analysis.

### 2.4. Overexpression and purification of recombinant K0513 protein

The generated plasmid construct pQE60-K0513 coding for the expression of K0513 protein was transformed into chemically competent *E. coli* XL1-Blue cells (a kind gift from Prof. A. Shonhai). Recombinant K0513 protein was expressed and purified using a previously described protocol [20]. The purification of the 6 x Histidine C-terminally tagged recombinant K0513 protein was conducted using nickel-affinity chromatography under native conditions. The recombinant protein was dialyzed overnight using an Amicon Ultra 15 mL spin dialysis column 10 000 MWCO (Merck, Germany). Purified protein was quantified using Bradford’s assay. Protein expression and purification was evaluated by SDS-PAGE analysis. The presence of recombinant K0513 was confirmed by Western blotting using mouse raised monoclonal anti-His antibodies (1 : 5000 dilution), and goat raised horseradish peroxidase (HRP) conjugated anti-mouse IgG antibodies (1 : 5000 dilution) [ThermoFisher Scientific]. The presence of K0513 was validated using polyclonal peptide specific anti-K0513 antibodies (1 : 500 dilution) [Abcam, UK], and HRP-conjugated goat anti-rabbit IgG secondary antibodies (1 : 3000 dilution) [ThermoFisher Scientific]. Peptide specific antibody against K0513, with N-terminal peptide segment from residue 7 - 123, (VGSLIDFGPEAPTSSPLEAPPPVLQDGDGSLGDGASESETTESADSENDMGESPSHPSWDQDRRSSSNESFSSNQSTESTQDEETLALRDFMRGYVEKIFSGGEDLDQEEKAKFGEY), was purchased from Abcam (AB121430, Abcam, Cambridge, United Kingdom). The presence of the secondary antibody was confirmed using the Pierce™ECL Plus Western blotting substrate (ThermoScientific, USA) and viewed using the ChemiDoc (BioRad, USA).

### 2.5. Tertiary structure analysis of recombinant K0513 protein

#### 2.5.1 Tryptophan-based fluorescence spectroscopy

The tertiary structural conformation of recombinant K0513 protein was analysed using tryptophan-based fluorescence spectroscopy as previously described [21]. Briefly, varying concentrations (0,1 - 0,5 µM) of recombinant K0513 protein was suspended in Tris-Buffered Saline (TBS: 50 mM Tris-HCl [pH 7,6], 150 mM NaCl) and incubated for 30 minutes at 20 °C. Fluorescence spectra were monitored using the JASCO-FP-850 Spectrofluorometer (Jasco, UK). The fluorescence emission spectra were monitored after initial excitation at 295 nm between 280 nm to 500 nm, at a scan speed of 1000 nm/min on a 10 nm emission bandwidth. The spectral measurements were averaged for three accumulations after baseline subtraction of buffer without protein to remove background emissions. In order to investigate the effect of nucleotides on the tertiary structure of K0513, the assay was repeated in the presence of different concentrations of nucleotides (0,2 – 1,0 mM) of GDP/GTP or (1,0 – 5,0 mM) ATP/ADP.

#### 2.5.2 Limited proteolysis

The conformational changes of K0513 that was expressed in *E. coli* XL1-Blue cells in the absence and presence of nucleotides using limited proteolysis was investigated. The assay was conducted as previously described [21]. Purified K0513 (6,42 μM) was incubated with 3,21 nM trypsin, using an enzyme to substrate ratio of 1:2000, at 37°C for 160 minutes, in the absence or of 5 mM ATP/ADP or 1 mM GTP/GDP. Proteolytic digestion of K0513 was analysed using 12 % SDS-PAGE.

### 2.7. Cell culture and Western blotting

All cell lines except the Human retinal pigment epithelial (RPE-1) were cultured and maintained at 37 °C in a humidified atmosphere of 5 % CO2 in air in Dulbecco’s Modified Eagle’s Medium (DMEM) + GlutMAXTM-1 containing 10 % Fetal calf serum (FCS) and 1% Penicillin-Streptomycin (10 000 U/ml) (Gibco Life Technologies, Grand Island, NY, USA). The RPE-1 cell line was cultured with 50 %DMEM/50% F10 Nutrient Medium supplemented with 10 % FBS, 0.01 mg/ml Hygromycin B and 1 % Penicillin-Streptomycin.

Whole cell lysates from all cell lines were prepared using a modified method described by Holden and Horton, 2009 with enhance radioimmunoprecipitation buffer [E-RIPA, 50 mM HEPES pH 7.4, 150 mM NaCl, 0.5 % Sodium deoxycholate, 1 % Sodium dodecylsulfate (SDS), 100 mM Dithiothreitol (DTT), 1U/ml Benzonase (Sigma), 1 mM EDTA, 1 mM PMSF and 0.1 % protease inhibitor cocktail (Roche Applied Science, Germany)]. The nutrient medium on cells was aspirated, washed twice with ice-cold phosphate buffered saline (PBS) pH 7.4 and lysed in E-RIPA buffer directly on the cell culture plate for 1 h at 4 °C [22]. The total protein content of each cell lysates was estimated using the BCA protein quantification kit (Pierce, Thermoscientific, USA). An equal amount of protein concentration was resolved on an SDS-PAGE in triplicates per cell line and electroblotted onto a nitrocellulose membrane. Immunoprobing of the blots were performed using anti-K0513 (AB121430, Abcam, Cambridge, United Kingdom) and GAPDH (endogenous control) antibody (14C10, Cell signaling, Danvers, USA). Chemiluminescence signals were visualized with the Clarity MaxTM Western ECL substrate (Bio-Rad, Italy) on the Azure c400 Geldoc system (Azure Biosystems, Inc, Dublin, USA).

## 3. RESULTS

### 3.1. K0513 is predicted to contain an SBF2 domain and a transmembrane region

Based on bioinformatics predictions, there are four variant transcripts of gene *K0513*, (variants 1 – 4; S1 Table) and two predicted variants (variants X1 and X2). *K0513* variants 1 and 2 code for the longer K0513 protein isoform a [K0513a; variant 2 differs in the 5’ untranslated region (UTR) compared to variant 1]. *K0513* variant 3 codes for isoform b, which is shorter but has the same N- and C-termini as isoform a (K0513b; variant 3 differs in the 5’ UTR and lacks an alternate in-frame exon compared to variant 1). *K0513* variant 4 codes for isoform c, which is shorter than isoform a and has a distinct C-terminus (K0513c; variant 4 differs in both the 5’ and 3’ UTRs and has multiple coding region differences, compared to variant 1). The human K0513a protein sequence is well conserved, and is found in eight homologues and 320 orthologues, based on data from NCBI. Phylogenetic tree analysis was used to establish orthology between the human K0513a and K0513 from eukaryotes including primates, rodents, ungulates, bony fishes and bats, and to compare their phylogenetic relationships (Fig 1). To the best of our knowledge, none of the K0513 protein homologues nor orthologues have been characterized. Not surprisingly, human K0513 clustered together phylogenetically with K0513 from primates, forming a monophyletic clade with chimpanzee K0513 (*Pan troglodytes*). Human K0513a was also found to cluster closely with K0513 from rodents and placentals (Fig 1).

K0513 is reported to contain a conserved SBF2 domain based on data obtained from UniProtKB, which is also reportedly present in about 52 % of the 320 orthologues of K0513. The SBF2 domain is found in Myotubularin related protein 13 (MTMR13; also called SBF2/SET binding factor 2), and is approximately 220 amino acids in length, based on data obtained from PFam (pfam12335) [23]. The significance of the SBF domain family (pfam12335) lies in its association with the Differentially Expressed in Normal and Neoplastic cells (DENN) domains, which is typically comprised of three parts. The original DENN domain flanked by two divergent domains, the uDENN (upstream DENN; pfam03456), and dDENN (downstream DENN; pfam03455). Some DENN domain containing proteins have been reported to interact with GTPases from the Rab family, functioning as nucleotide exchange factors [24], and others have been reported to be involved in regulation of Mitogen-activated protein kinases (MAPKs) signalling pathways. The myotubularin family is a highly conserved group of ubiquitously expressed phosphatidylinositol 3-phosphatases [25]. The 208 kDa MTMR13 protein is composed of various domains including the tripartite DENN domain, an SBF2 domain, a glucosyltransferase/Rab-like GTPase activator/myotubularin (GRAM) domain, a pseudo-phosphatase domain and a pleckstrin homology (PH) domain. PH domains are involved in membrane association and usually bind phosphates [26,27,28]. The GRAM domain is known to mediate membrane attachment by binding to phosphoinositides [28]. MTMR13 also forms part of the 18 known DENN domain proteins encoded in the human genome [29].

A multiple sequence alignment of K0513a, K0513b, K0513c and SBF2-containing protein, MTMR13, showed that the K0513 protein sequence is associated with the dDENN and SBF2 domains of MTMR13 (Figs 2 and S1). K0513 is partially aligned to the cDENN domain of MTMR13 (Figs 2 and S1). MTMR13 has previously been reported to facilitate exchange of nucleotides for Rab21 and Rab28 [30]. Rab (Ras-related protein in brain) proteins represent the largest family of the small GTPases, with over 70 members in humans [31]. Rab GTPases are important regulators of intracellular membrane trafficing in all eukaryotes [32]. These GTPases are known to interact with effectors such as kinases, phosphatases and tubular-vesicular cargo [33]. Rab GTPases localize to certain intracellular membranes and recruit various Rab effectors including molecular motors, tethering factors and phospholipid modulators [34,35]. They have been described as “switch molecules” in their ability to alter between an active GTP-bound state and an inactive GDP-bound state. In its active state, the Rab GTPase will attach to the membrane and regulate its trafficking through secretory and endocytic pathways, which is accomplished through interaction with effector proteins known for the connection of activated Rab proteins to a diversity of cellular events such as cytoskeletal dynamics, phosphatidylinositol signaling events and protein kinase-mediated signal transduction [36,37]. For Rab to localize to the membrane, prenylation at the C-terminal cysteine residues is required, and the only known function of prenylation is to confer a hydrophobic character onto a cytosolic protein, thus giving the recipient protein the physical capacity to make a steady attachment with a lipid bilayer [36]. During this process, Rab binds the Rab escort protein (REP), after which Rab geranylgeranyltransferase prenylates the Rab GTPase, which is then transported to the target membrane [38]. Rab is then activated for downstream signalling by a guanine exchange factor (GEF), which facilitates the exchange of guanine-5’-diphosphate (GDP) for guanine-5’-triphosphate (GTP) [35]. Whilst activated, Rab associates with various effector proteins that facilitate the movement of vesicles from donor membranes to target membranes [39]. Rab is then inactivated by binding GTPase activating proteins (GAPs), which hydrolyse GTP to GDP. Finally, inactive Rab is removed from the membrane by the GDP dissociation inhibitor (GDI), which solubilizes the GTPase, thereby preparing Rab for the next cycle of vesicular transport [40].

**Fig 2.**
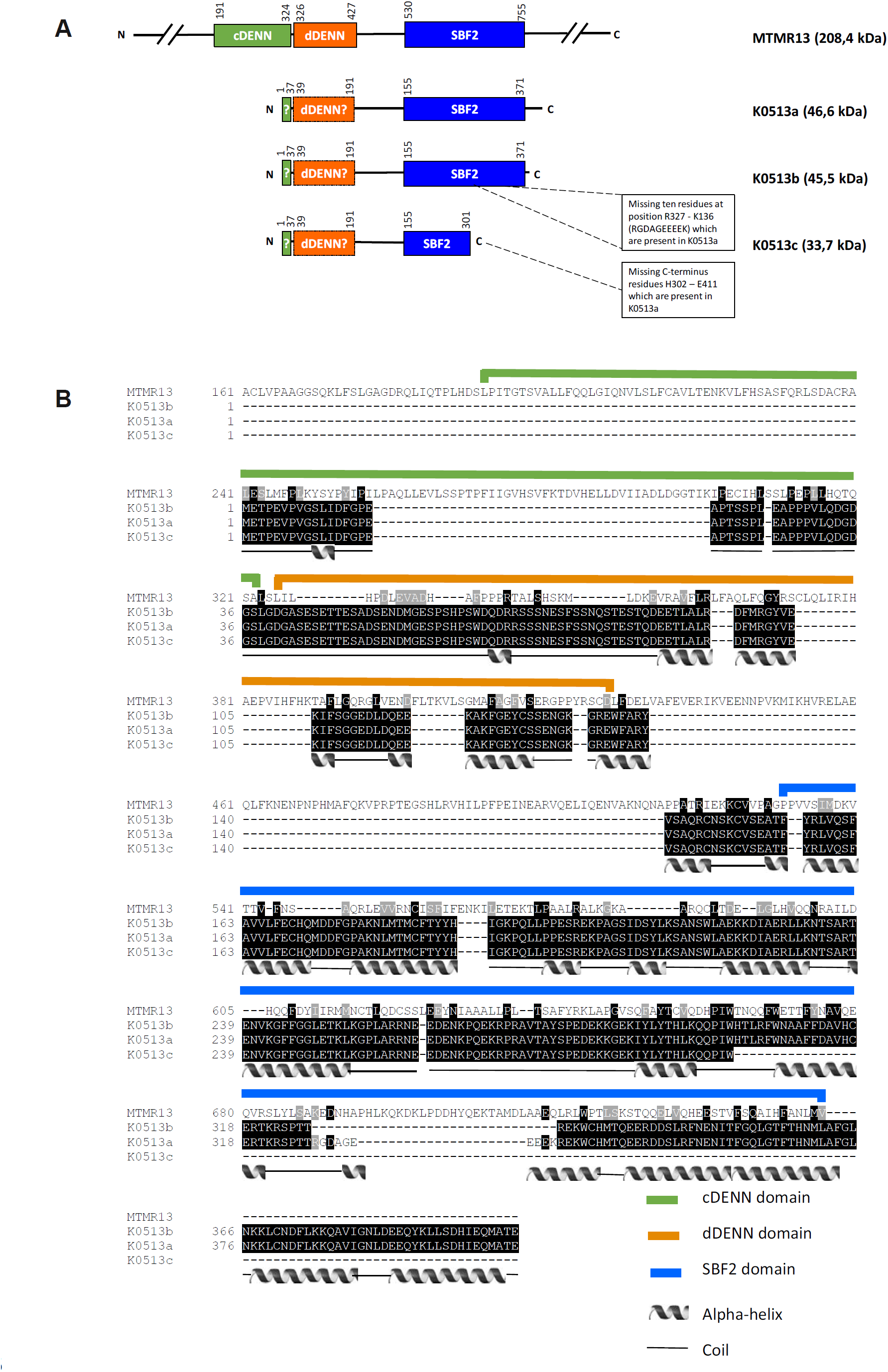
Schematic of the domain organization and multiple sequence alignment of K0513 isoforms and MTMR13 DENN and SBF2 domains. K0513 was aligned with the known DENN domain containing GEF from the myotubularin family, MTMR13. Segments highlighted by green, orange and blue lines represent domains found in MTMR13 and their respective positions. The hypothetical positions of the cDENN, dDENN and SBF2 domains are highlighted by olive green, orange and blue, respectively. Prediction of the secondary structures of human K0513a was performed using PROMALS3D [13], where helices represent alpha helices, and black lines represent coils. The alignment was generated using Clustal Omega [12].

The DENN domain is found in a variety of signalling proteins involved in Rab-mediated processes or regulation of mitogen-activated protein kinase (MAPK) signalling pathways [24]. DENN domain proteins are GEFs for specific Rab proteins [30,41,42]. Studies have shown that the DENN domain by itself is adequate for guanine exchange function (GEF) activity [41,42]. MTMR13 promotes the exchange of GDP for GTP in the small GTPase proteins, Rab21 and Rab28 [29,43]. MTMR13 converts the inactive GDP-bound Rab28 into its active form, GTP-bound Rab28 protein [30]. The conserved SBF2 domain and cDENN domain in K0513 suggests that K0513 may potentially interact with a Rab protein, and secondly, may play a role in the nucleotide exchange of Rab proteins. The secondary structure of K0513 was predicted to be comprised of alpha helices and coils, with the helices taking up 70 % of the whole structure (Fig 2).

The amino acid composition and physiochemical properties determine the fundamental properties of a protein. K0513 has a molecular weight of 46,638 kDa and is composed of 411 residues; it has a theoretical isoelectric point of 4,7 (Fig 3). The primary structure analysis shows that K0513 is strongly dominated by polar residues, constituting 55.5 % of the protein thus making the protein hydrophilic in nature (Fig 3). Interestingly, the presence of several cysteine residues suggests that disulphide bridges may constitute a strong component regulating the folding of HsK0513, however, there are none predicted to occur based on DISULFIND (S2 Fig). The computed isoelectric point (pI) of human K0513 is 4.6994, suggesting that the molecule is acidic in nature. This is due in part to the high number of glutamic acid (Glu) residues (Fig 3). The secondary structure of human K0513 was further analysed for hydropathy, amphipacity and topology to predict whether K0513 has capability to bind to the cell membrane in relation to its role as a GEF for Rab, studies have shown that some GEFs are recruited to specific organelle membranes via their transmembrane regions, thereby localizing the GTP-bound Rab to the same membrane [43]. Residue 150 to 167 were predicted to be hydrophobic with the hydrophobicity score of 1.2 and the region is suggested to form part of the transmembrane anchor (Fig 3). Both the N-terminal and the C-terminal are predicated to be cytosolic and anchored to the membrane through the transmembrane region (Fig 3).

**Fig 3.**
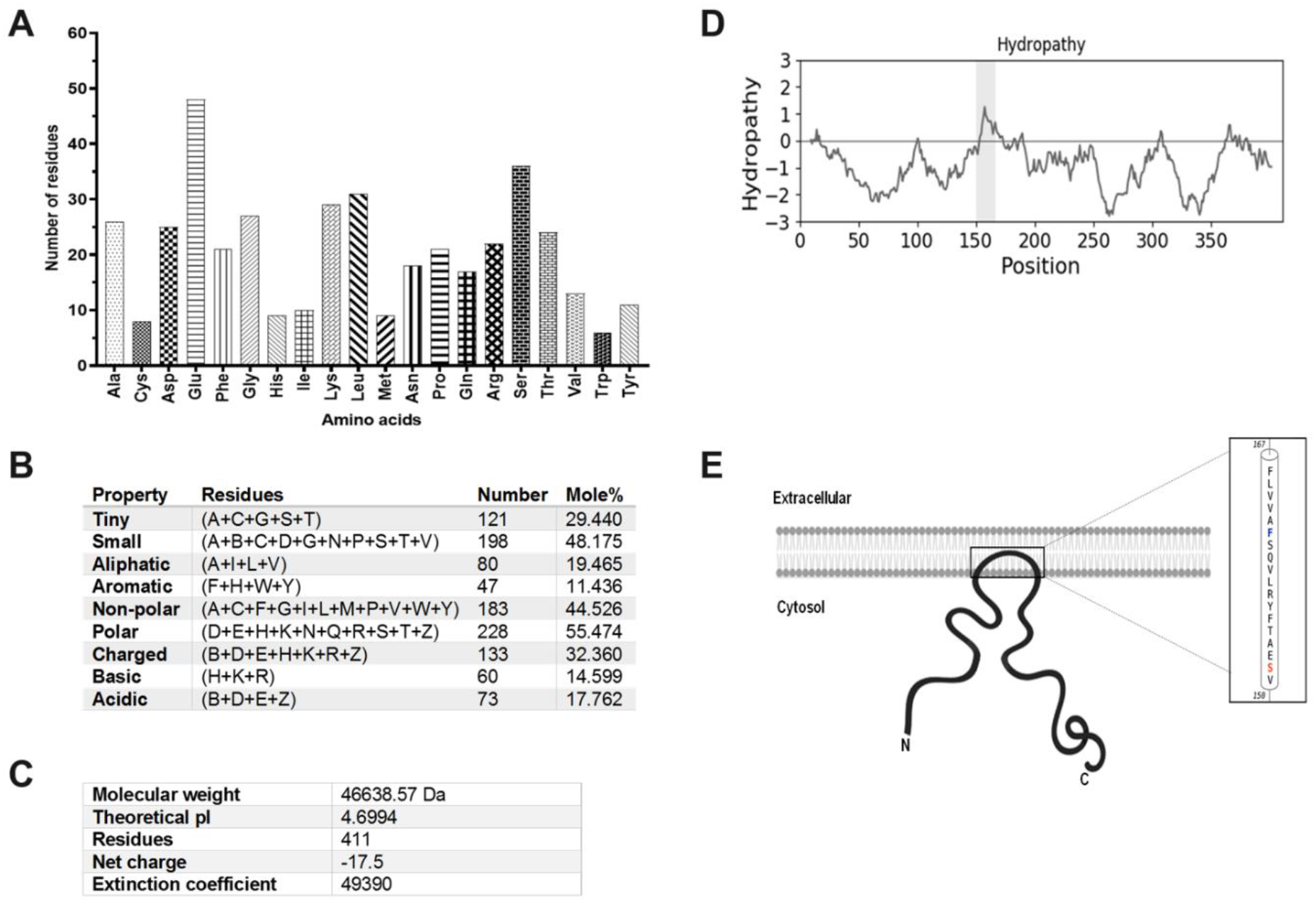
Amino acid composition and physicochemical properties HsK0513. The amino acid composition of K0513 was computed using PEPSTAT module integrated in the EMBOSS software (A). The amino acid properties of amino acids are summarized in (B). Physicochemical characterization of the protein was computed using the Expasy’s ProtParam server (C). Hydropathy plot showing the degree of hydrophobicity and hydrophilicity, the hydrophobic transmembrane region is highlighted in grey (D). The Hidden Markov Model for Topology Prediction (HMMTOP) showing helical-transmembrane residues and the position of the carboxyl-terminal and amino-terminal relative to the cell cytosol (E). The models were generated using Biotools Web-based Hydropathy, Amphipathicity and Topology (WHAT) [15].

### 3.2. The SBF2 domain of K0513 appears to exhibit a coiled coil fold

The predicted three-dimensional structure of human K0513 exhibits a globular fold and the N-terminal loop is positioned towards the C-terminus (Fig 4A). The N-terminal and the C-terminal regions of K0513 are more hydrophobic as compared to the central region of the protein (Fig 4B). The K0513 homology model was generated from several templates using Phyre^2^, for which the highest probability (58,1 %) for shared homology was the Glutathione S-transferase C-terminal domain-like template (Fig 4). The SBF2 domain of K0513 resembles a coiled coil (Fig 4A). GRAB protein has been shown to function as a physiological GEF for Rab3a, specifically through its coiled coil domain [44].

**Fig 4.**
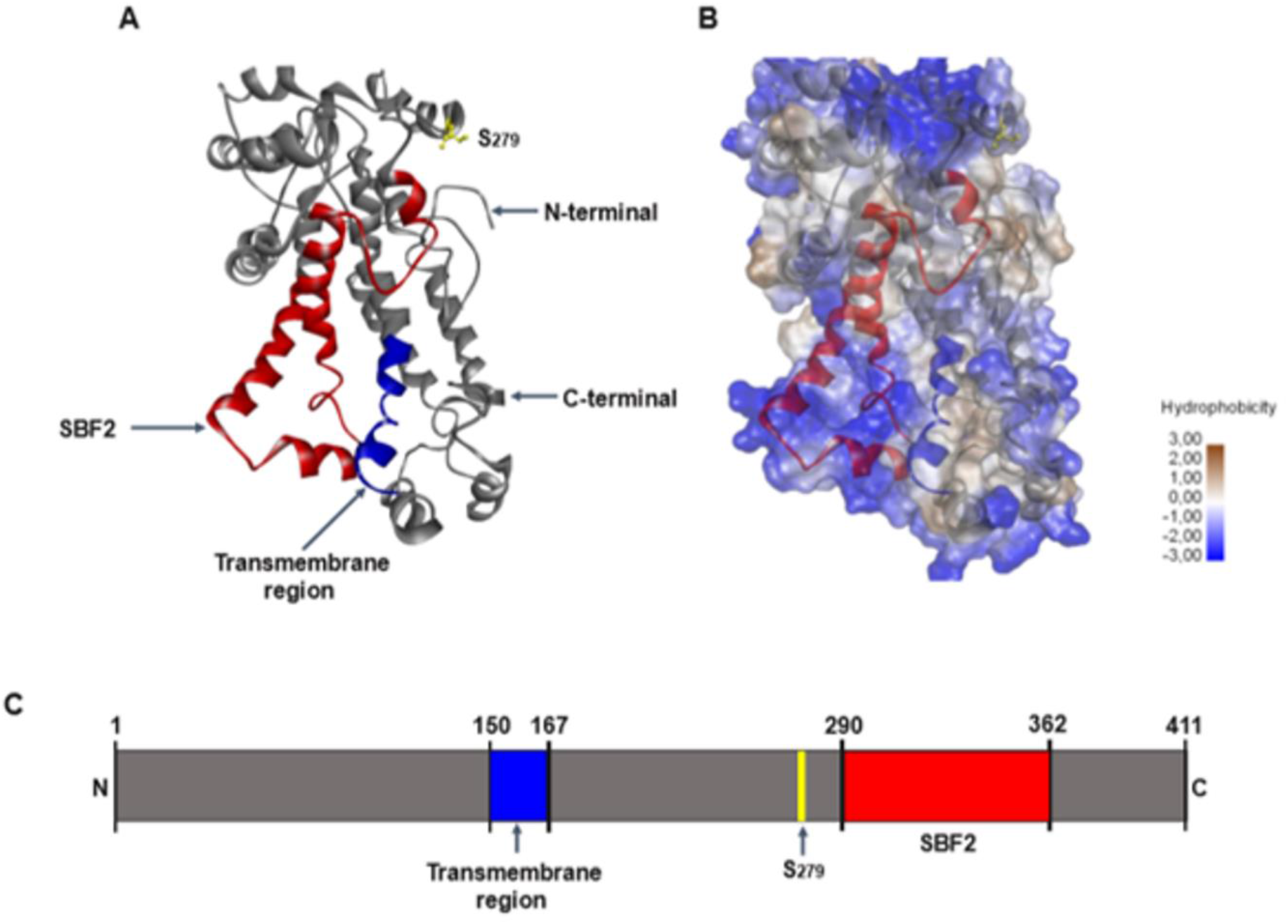
Three-dimensional homology model of K0513. The 3D model of K0513 showing the SBF2 domain (yellow) and the phosphoserine residue represented in balls and sticks (pink) (A). The carboxyl-terminal is represented by red whereas the amino-terminal is represented with blue colour. The hydrophobicity surface plot showing hydrophobic (brown), amphipathic (grey) and hydrophilic residues (blue) (B); as well as the schematic presentation of KO513 structure (C) are shown. The model was generated by Phyre^2^ [14] and visualized using Biovia Discovery studio [18].

Considering that the three-dimensional structure of a protein may yield valuable information about its functional properties, the structural similarity between proteins may be a good indicator of functional similarity. Using Phyre^2^, some interesting proteins were predicted to partially align with regions of K0513 (S2 Table). Amongst these was lipid binding protein (PDB ID: c2jobA), which are normally molecules that may reversibly and non-covalently associate with lipids, and are often membrane-associated [45]. Another pair of interesting templates are regulator of g-protein signalling 7 (RGS7; PDB ID: c2d9jA) and RGS2 (PDB ID: c2af0A), which both act as G-protein activating proteins (GAPs) interacting directly with the activated G-protein and whose function is to hydrolyse GTP which in turn promotes inactivation of the G-protein (S2 Table) [46]. Could K0513’s function possibly be related to these proteins, with respect to association with cellular membrane and facilitating nucleotide exchange of GTPases?

### 3.3. Predicted interactome of human K0513

The protein-protein interaction prediction revealed 71 possible interactors of K0513 (S3 Table), interaction partners were generated through the online Integrated Interaction Database (IID) prediction. The prediction identified molecules that had previously been established to interact with K0513 through the yeast two-hybrid system, thereby validating the predictions through experimental data (Table 3) [6]. These included INTS4, which is involved in the small nuclear RNAs (snRNA) U1 and U2 transcription; HAX1, which regulates reorganization of the cortical actin cytoskeleton, and KIBRA/WWC1, which plays a pivotal role in tumour suppression by restricting proliferation and promoting apoptosis (Table 3) [6]. The predicted interactome included various proteins implicated in GTPase activity. The most interesting of these was predicted interaction of Rab3a, a small GTPase that selectively binds secretory vesicles and switches between the active vesicle associated GTP-bound form and the inactive cytosolic GDP-bound form (Fig 5A; S3 Table) [47]. Another predicted interactor of K0513 is Syntaxin-binding protein 1 (STXBP1, also called MUNC18-1) [47], which is involved in the regulation of synaptic vesicle docking and fusion through specifically binding with GTPases (Fig 5A). Interestingly, Rab3A has been reported to interact directly with Munc18-1 in vesicle priming during exocytosis (Fig 5A; S3 Table) [47]. The protein, IQ motif and SEC7 domain-containing protein 3 (IQSEC3), is also an interesting predicted interactor of K0513, in particular in light of its role as a guanine exchange factor for the small GTPase ADP-ribosylation factor (Arf6), which plays a role in intracellular vesicular trafficking, and regulates the recycling of various types of cargo (Fig 5A; S3 Table) [48]. The last predicted interactor of K0513 involved with GTPase activities is Rho GDP-dissociation inhibitor 3 (ARHGDIG), which functions to inhibit the GDP/GTP exchange reaction of RhoB; RhoB is a small GTPase that regulates vesicle transport and actin reorganization (Fig 5A; S3 Table) [49].

**Fig 5.**
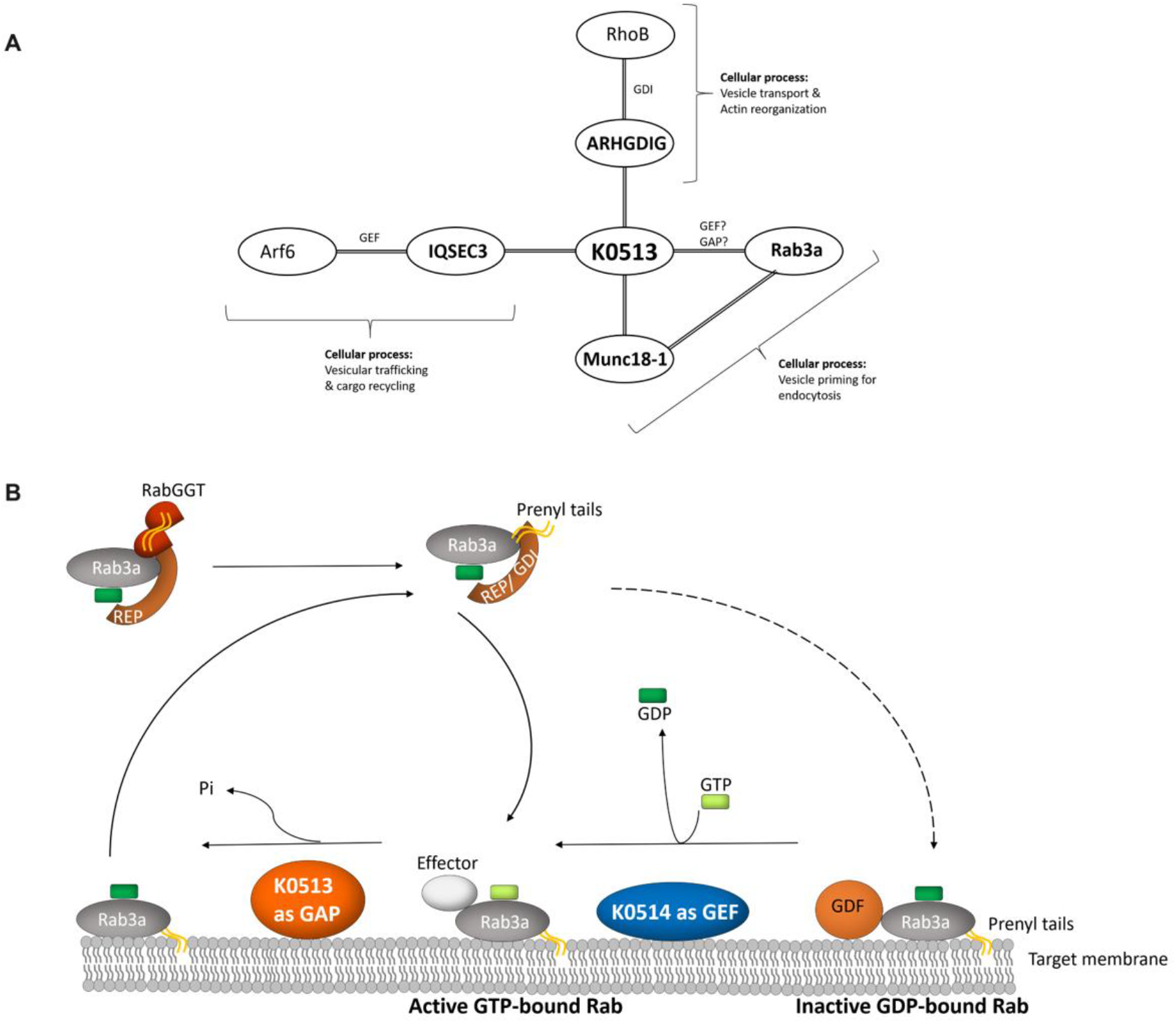
Predicted interactome of K0513 and proposed roles of K0513 in the Rab3a signal transduction pathway. (A) K0513 is predicted to interact with Rab3a, Mun18-1, and Munc18-1 has been shown to interact with Rab3a in processes facilitating vesicle priming for endocytosis; K0513 is predicted to interact with IQSEC3 which functions as a GEF for Arf6, which facilitates vesicular trafficking and cargo recycling; K0513 is predicted to interact with ARHGDIG1 which is the GDP dissociation inhibitor for RhoB, and facilitates vesicular transport and actin reorganization. (B) Newly synthesized Rab interacts with Rab escort protein (REP) in its GDP bound state, recruiting the small GTPase to Rab geranylgeranyl transferase (RabGGT) which prenylates Rab at C-terminal cysteine residues. Rab is then activated by a guanine exchange factor (GEF) which exchanges GDP for GTP and facilitates recruiting Rab to specific cellular membranes. GTP-bound Rab binds effector proteins that mediate various steps in vesicular trafficking. 2. Rab is inactivated by interacting with GTPase activating protein (GAP) through hydrolysis of GTP to yield GDP-bound Rab. 3. Prenylated Rab is removed from the membrane and solubilized by interacting with GDP dissociation inhibitor (GDI), which shields the nonpolar geranylgeranyl groups in the prenyl tails from the polar environment. 4. GDI dissociation factor (GDI) separates Rab from GDI and so allows Rab to be inserted into the target membrane, in preparation for the next cycle.

Our *in silico* analysis of human K0513 would suggest the association of K0513 with Rab GTPases. We have seen that K0513 contains a conserved SBF2 domain, and a partially conserved DENN domain (Fig 2), K0513 exhibits regions of sequence similarity to proteins involved in regulation of GTPases (S2 Table), and is predicted to interact with a Rab GTPase, and proteins involved in their regulation (S3 Table). However, it is still unclear in what capacity K0513 may interact with the Rab GTPase. Whether K0513 hydrolyses GTP, thereby acting as a GAP protein; or dissociates a GAP from Rab GTPase, thereby acting as a GDF; or even directly associates with Rab GTPase, inducing a conformational change which facilitates the exchange of GTP for GDP, and thereby acting as a GEF, is unclear and remains to be determined (Fig 5). We were therefore curious to investigate whether nucleotides could induce conformational changes K0513 through biophysical analysis to develop a better understanding.

### 3.4. Overexpression and purification of recombinant K0513

The *E. coli* expressed recombinant K0513 protein migrated as a species of 47 kDa on SDS-PAGE, which correlated to the predicted molecular weight. K0513 protein was purified in its the native form using Nickel-affinity chromatography (Fig 6). The authenticity of the successfully expressed and purified protein was validated by Western blotting target specific antibodies (Fig 6). The overall purification yielded 20 mg per 1 L total culture volume of approximately 80 % purity (Fig 6).

**Fig 6.**
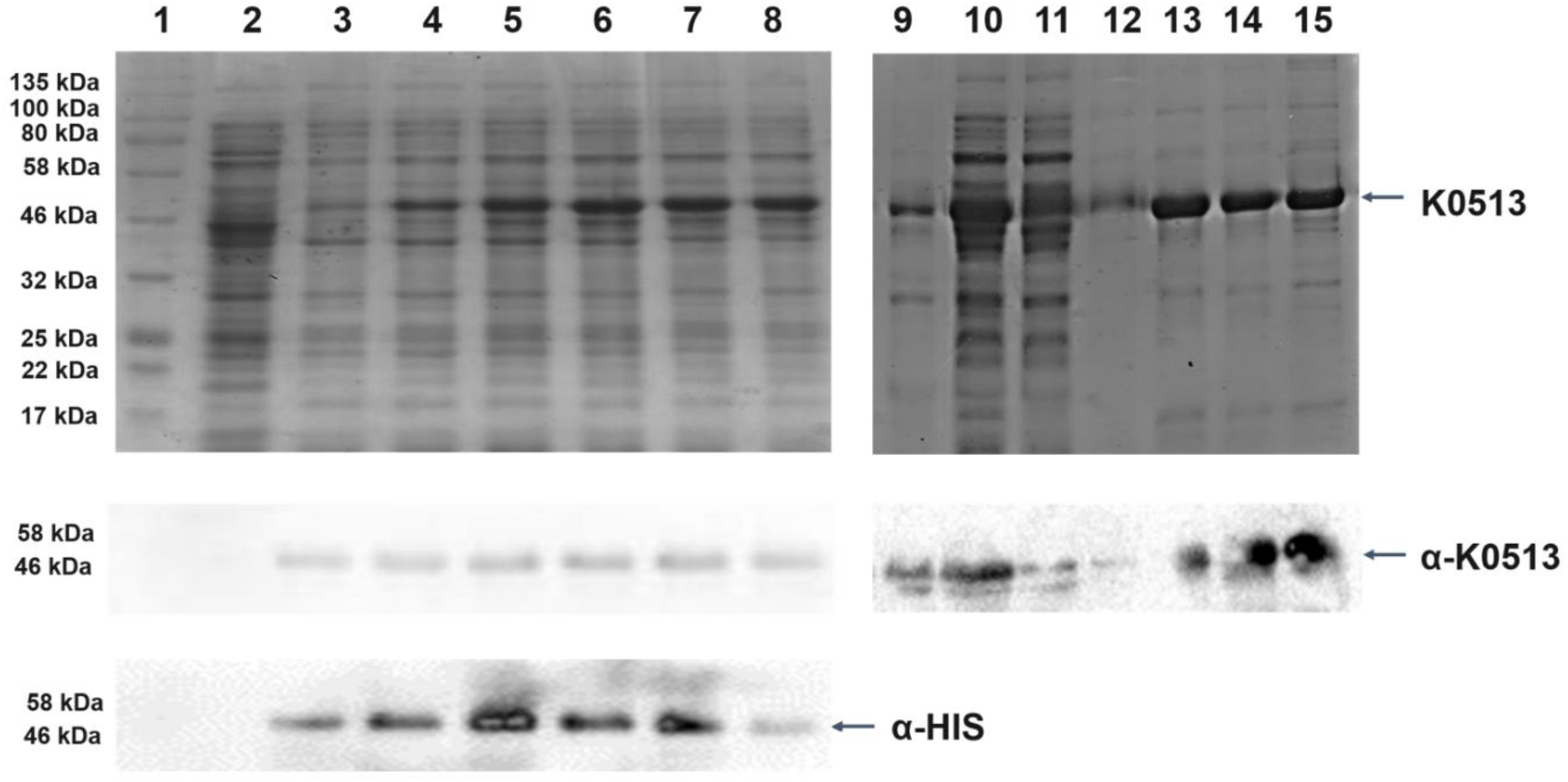
Heterologous expression and purification of recombinant K0513 using nickel affinity chromatography. SDS-PAGE (12 %) analysis of K0513 expression in *E. coli* XL1-Blue cells. Lanes 1 – Protein marker; 2 – total extract of cells transformed with pQE60 as control; 3 – total extract of cells transformed with pQE60-KIAA prior to IPTG induction; 4 - 8 – total cell lysates obtained up to 5 hours post IPTG induction; 9, 10 – pellet and soluble fractions obtained from total lysate of cells transformed with pQE60-K0513, respectively; 11 - flow through fraction; 12 – wash fraction, 13 - 15 – elution fractions. Western blotting was performed using anti-K0513 and Anti-Histidine antibodies.

### 3.5. Investigation of the effect of nucleotides on the tertiary structure of K0513

We investigated the effects of nucleotides on the tertiary structure of the recombinant K0513 protein using tryptophan-based fluorescence assay. The K0513 protein sequence possesses a total of six tryptophan residues at positions W65, W134, W218, W300, W306 and W339. The conformation of K0513 in the presence of nucleotides ATP/ADP and GTP/GDP was assessed (Fig 7). Recombinant K0513 had a maxima of 332 nm at 0 mM ATP, ADP, GTP and GDP. We observed a marginal red shift in the emission maxima of recombinant K0513 exposed to increasing concentrations of ATP, ADP, GTP and GDP of approximately 3 nm, 3 nm, 4 nm and 2 nm, respectively (Fig 7). In addition, K0513 was incubated with a fixed concentration of GTP and the fluorescence was monitored over time (Fig 7). There was no significant red or blue shift of the 332 nm peak. This data signifies that the conformation of K0513 remains unaltered in the presence of nucleotides, which would suggest that K0513 is not binding nor hydrolysing nucleotides.

**Fig 7.**
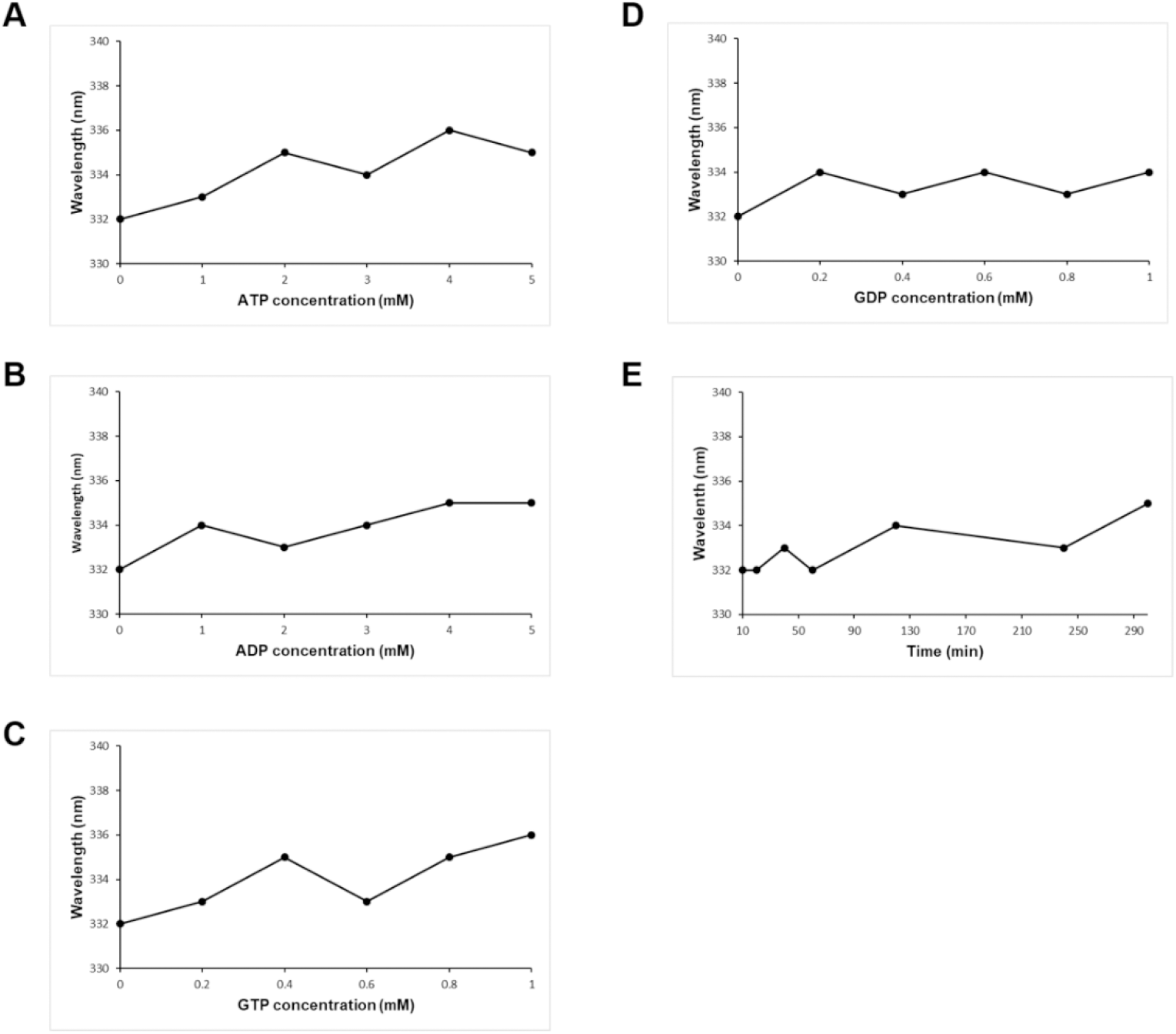
Effect of nucleotides on the tertiary structure of K0513. The conformational change of K0513 was assessed in the presence of various concentrations of ATP and the shift in wavelength was plotted against concentration (A), the same was followed for ADP (B), GTP (C) and GDP (D). The conformational change of K0513 was assessed in the presence of a fixed concentration of GTP at different time points, and the shift in wavelength was plotted against time point in minutes (E). Initial excitation was at 295 nm and the spectra were monitored between 280 to 500 nm, 1000 nm/min scan speed, 3 accumulations and 10 nm emission bandwidth.

To determine whether the observed fluorescence spectral shifts were due to conformational changes in the protein tertiary structure, chaotropic denaturants were used. The maximum fluorescence intensity for native K0513 was observed at 332 nm, which is expected [50]. This would suggest that K0513 assumes a folded state prior to introduction of the denaturants. The addition of urea and guanidine HCl to recombinant K0513 at increasing concentrations resulted in a significant red shift for both denaturants (Fig 8). The red shift signifies that recombinant K0513 is now in a more unfolded and extended state due to resulting surface exposed charged residues [51]. Taken together, this data would suggest that K0513 conformation is not perturbed by the presence of nucleotides.

**Fig 8.**
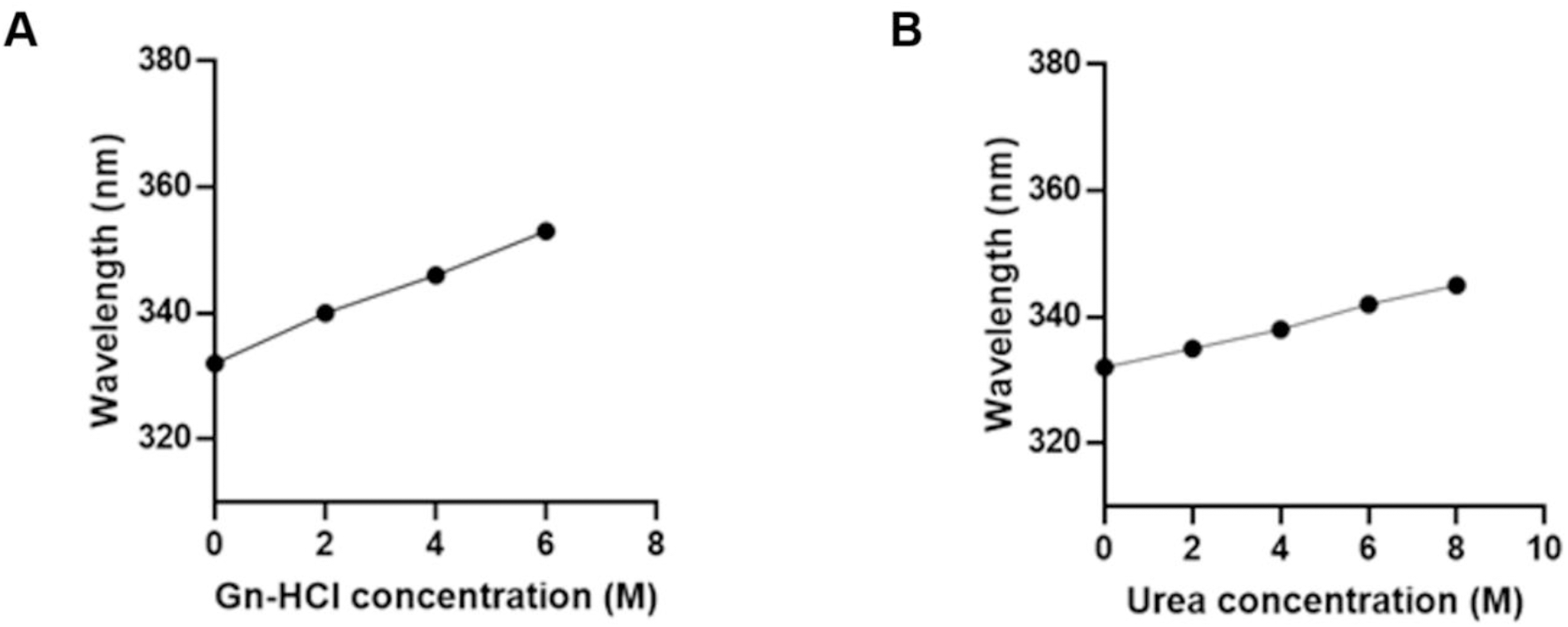
Conformation of K0513 in the presence of guanidine-HCl and urea. K0513 conformation was assessed using tryptophan fluorescence spectrophotometry in the presence of various guanidine-HCl concentrations, and the wavelength shift was plotted against its concentration (A); the same was followed in the presence of urea (B). Initial excitation was at 295 nm and the spectra were monitored between 280 to 500 nm, 1000 nm/min scan speed, 3 accumulations and 10 nm emission bandwidth.

To further validate the results that were obtained using the tryptophan-based fluorescence spectroscopy, we performed a trypsin based limited proteolysis assay in which we monitored the effect of nucleotides on the conformation of recombinant K0513 (Fig 9). The protein was digested to smaller fragments, as was represented by smaller bands on SDS-PAGE. This demonstrated that recombinant K0513 is a proteolytically susceptible molecule. The fragmentation profiles obtained by limited proteolysis of K0513 in the presence of various nucleotides ATP/ADP and GTP/GDP were not distinct (Fig 9). This suggests that the various nucleotides did not uniquely modulate the structure of the protein. This is in support of the data obtained by intrinsic fluorescence analysis (Fig 9).

**Fig 9.**
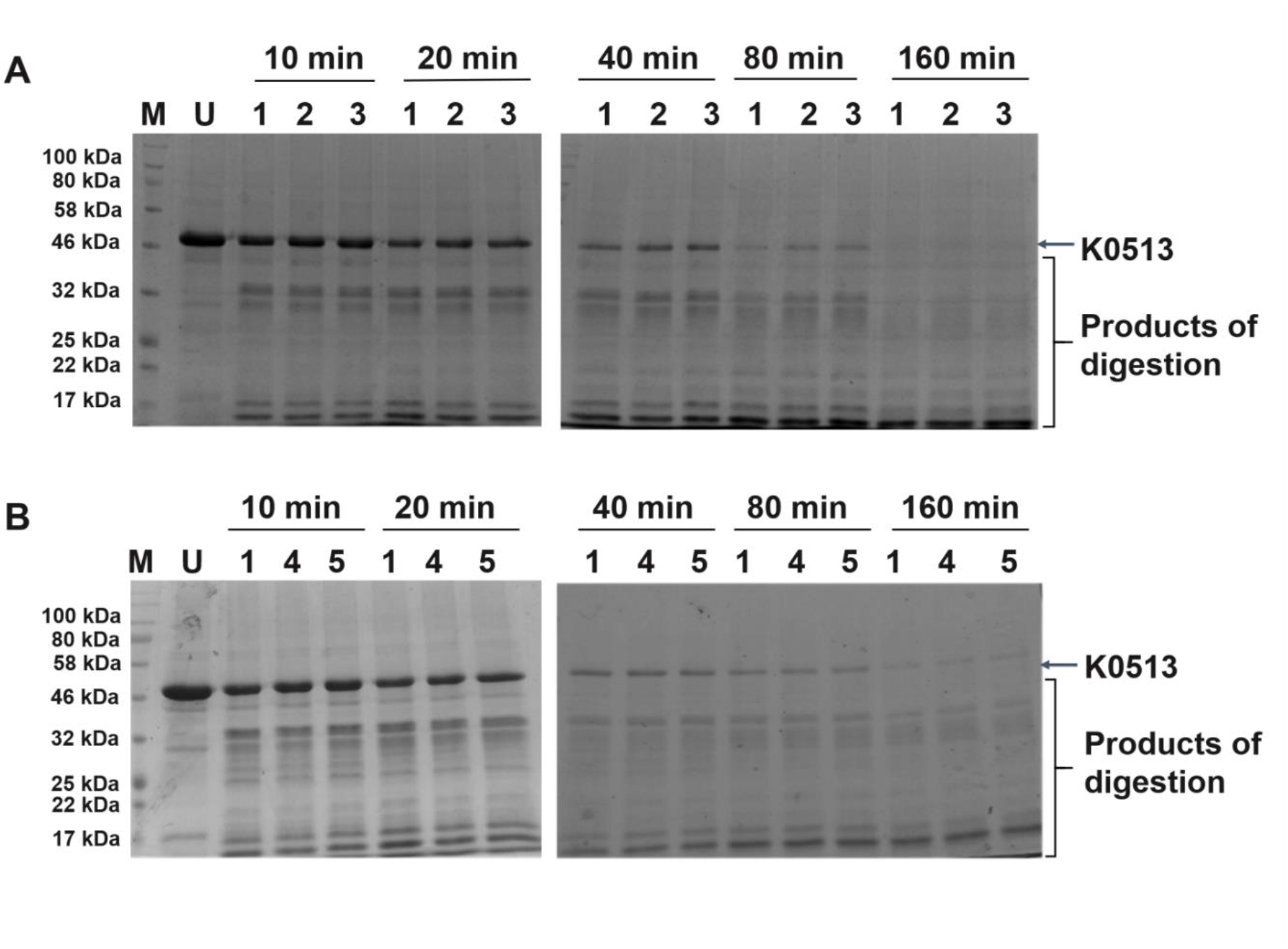
Proteolytic digest of K0513 in the absence and presence of nucleotides. SDS-PAGE analysis of recombinant K0513 that was subjected to limited proteolysis (LP) in the presence of ATP/ADP (A) and in the presence of GTP/GDP (B). Lanes M -Protein marker; U – undigested K0513; 1 - K0513 subjected to LP; 2 – K0513 in the presence of 5 mM ATP subjected to LP; 3 - K0513 in the presence of 5 mM ADP subjected to LP; 4 - K0513 in the presence of 1 mM GTP subjected to LP; 5 - K0513 in the presence of 1 mM GDP subjected to LP. The limited proteolysis by trypsin was conducted using an enzyme to substrate ratio 1 : 2000. The duration of exposure to trypsin is given in minutes.

We further investigated the presence of K0513 in various lysates prepared from both normal and cancerous human cell lines. Using Western blotting, K0513 was detected in all the cell lines tested including retinal pigment epithelial cells, keratinocytes, lung carcinoma cells, fibrosarcoma cells, and cervical adenocarcinoma cells, albeit at very low levels in the latter (Fig 10). Notably, K0513 was detected as a doublet band in both the epithelial and fibrosarcoma lysates at ∼ 45 kDa and ∼ 46 kDa, which could signify the presence of two K0513 isoforms (Fig 10). This finding opens the door for further establishment of the role of K0513 in cancer cells.

**Fig 10.**
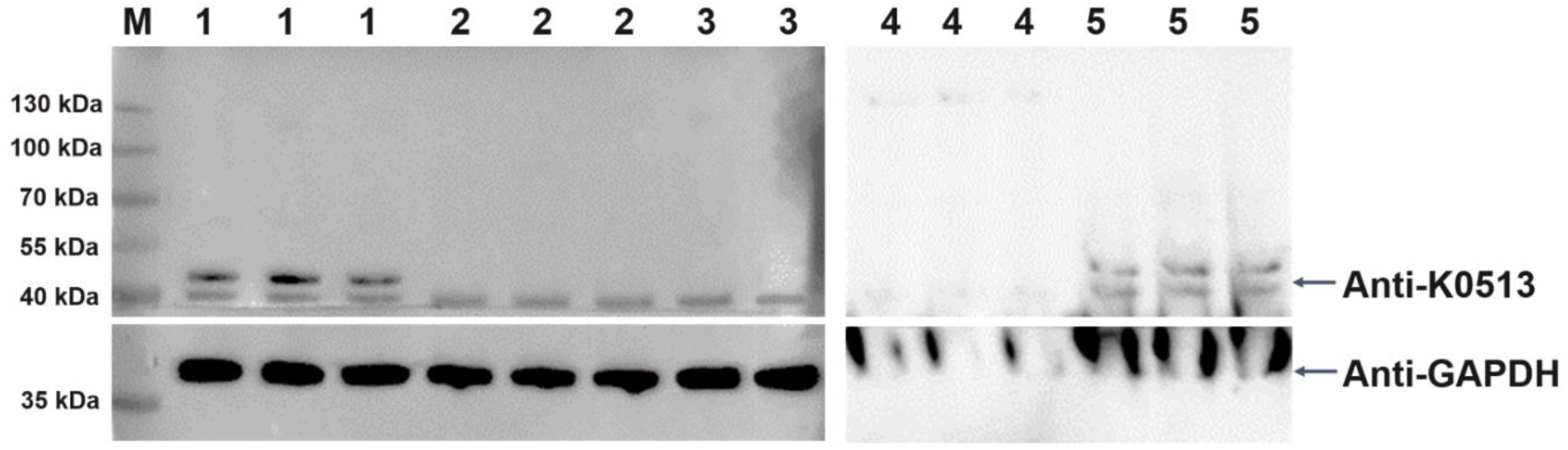
Detection of K0513 in human lysates prepared from different cell lines to assess the anti-K0513 antibody. K0513 was detected in various human cell lines by Western blotting using the rabbit raised anti-K0513 antibody at a 1:100 dilution. Lanes M – Protein marker; 1 - Human retinal pigment epithelial (RPE-1) whole cell lysate; 2 - Human immortalized keratinocyte (HaCaTs) whole cell lysate; 3 – Human lung carcinoma (A549) whole cell lysate; 4 - Human cervical adenocarcinoma (HeLa) whole cell lysate; 5 – Human fibrosarcoma whole cell lysate.

## 4. DISCUSSION

Our findings which confirmed that human K0513 contains conserved SBF2 and dDENN domains, and a transmembrane region, suggest a possible role of this protein as nucleotide exchange factor for Rab GTPases. *In silico* analysis revealed the protein to be predominantly composed of alpha helices and coils, and homology modelling showed the fold of K0513 to globular in nature. Investigating the predicted interactome of K0513 revealed its possible association with proteins related to GTPase pathways, in particular, Rab3a, a small GTPase protein. Furthermore, we optimised a protocol for the expression and purification of K0513. Biophysical characterisation of the protein established that the tertiary fold of K0513 is not modulated by nucleotides, suggesting that it may not directly interact with nucleotides, as observed using both tryptophan-based fluorescence and limited proteolysis assays. In addition, data obtained from the tryptophan-based fluorescence assay revealed a pronounced red shift of the protein in the presence of denaturants, suggesting that, in its native state, K0513 exists as a globular protein. This is the first report of recombinant K0513 protein production from a bacterial system, and analysis of tertiary structure. Considering that we found the conformation of K0513 to be unperturbed by nucleotides, together with the *in silico* analysis, we propose that K0513 may function as a GEF of Rab3a. Our findings further suggest that K0513 does not necessarily bind to G-proteins (GAP function) to accelerate their GTPase activity through hydrolysis of GTP.

Conservation of protein structure often mirrors similarities in protein function, and so this study sought to identify the structure-function features of “uncharacterized protein” K0513. K0513 comprises a conserved SBF2 domain, which is present in MTMR13. The SBF2 domain is just downstream of the dDENN domain (Fig 2). The DENN domain containing proteins are reported to be involved in a variety of signalling pathways, however, there is a significant degree of variation within these domains from protein to protein [29]. The DENN domain found in MTMR13 has been suggested to facilitate nucleotide exchange function of specific small GTPases from the Rab family [30]. The DENN domain binds to switches I and II of Rab21 GTPase resulting in conformational change in the region to allow GDP to GTP exchange [52]. The tripartite DENN module is comprised of the upstream DENN (uDENN), followed by central DENN (cDENN), and the downstream DENN (dDENN). The uDENN may occur independently of the DENN module and interact with various GTPases [53]. The central cDENN and dDENN occur together, the d-DENN is reported to make fewer contact with the GTPase thus it always associates with cDENN to maintain the GEF function [52]. A multiple protein sequence alignment of MTMR13 and the K0513 isoforms interestingly exhibited some similarity of K0513 over the cDENN and dDENN domains of MTMR13 as well as the SBF2 domain (Fig 2). From the sequence conservation observed between K0513 and the DENN and SBF2 domains of MTMR13, we asked whether protein K0513 cross-talks to a small GTPase protein, possibly through a nucleotide exchange function. There is no evidence on how the SBF2 domain facilitates the nucleotide exchange activity of the DENN domain, however, their co-existence strongly suggests they have a functional relationship. Homology determination is important in ascertaining functional features in protein structures especially in the absence of a crystal structure, as well as structural prediction of proteins. The query protein sequence can be aligned with identified homologues for determination of domains and motifs based on the level of conservation between the protein. Although K0513 is well conserved amongst its homologues, K0513 exhibits low conservation to proteins that have had their crystal structures resolved, and as a result the three-dimensional model of K0513 was generated using segments of different templates (Fig 4). The three-dimensional homology model of K0513 shows a globular fold, and the SBF2 domain appears to be surface exposed and a coiled coil in structure (Fig 4). The Phyre^2^ tool also showed segments of proteins that align to regions of K0513; it was notable that some of these proteins are associated with the GTPase pathway, in particular two GAP proteins (S2 Table).

Protein-protein interaction prediction is crucial in ascertaining pathways and cellular processes that the protein is involved in. This study used the Integrated Interaction Database (IID) to predict protein-protein interactions [19]. IID is the only database with context-specific networks for the common model organisms and domesticated species [19], it integrates experimentally detected protein interactions from nine curated databases (BioGRID, IntAct, I2D, MINT, InnateDB, DIP, HPRD, BIND, BCI). Predicting the K0513 interactome may reveal signalling pathways that K0513 could be involved in and so enable prediction of function. Physicochemical analysis showed that K0513 possess a transmembrane region. The 17 residues long alpha-helical region consists of hydrophobic residues positioned from amino acid residue 150 to 167. Transmembrane proteins can be integrally anchored to the lipid bilayer for interaction with membrane proteins. For example, general receptor for phosphoinositides 1 (Grp1), a known GEF interact with PI3-kinase in the inner membrane through its pleckstrin homology (PH) domain [54]. When anchored to the membrane, Grp1 catalyses the nucleotide exchange and activation of ADP-ribosylation factor-6 (Arf6) [55]. The presence of the transmembrane suggest that K0513 performs its nucleotide exchange function on membrane localised Rab GTPases when it is anchored to the lipid bilayer. Furthermore, it may directly or indirectly facilitate cellular processes such as vesicle trafficking or serve in transducing signal in pathways such as neuroplasticity, cytoskeletal regulation and apoptosis [6]. The predicted interaction with the transmembrane proteins, most notably the Na^+^/Ca^2+^ Solute Carrier Family 8 Member A2 (SLC8A2) further substantiate this finding. Most importantly, Rab3a, a GTPase involved in vesicle trafficking has been predicted to interact with K0513 supporting the role of K0513 along the membrane. Another important finding is the identification of the possible phosphorylation at S279. Protein phosphorylation serves as an initial step crucial for facilitating cellular and organic functions such as regulation of metabolism, subcellular trafficking, and signalling transduction [56]. Phosphoresidues can be activated or deactivated by kinases or phosphatases, this modification induces conformational change when the protein interacts with other proteins. Some GEFs require phosphorylation to activate their GEF activity, p115, the smallest member of the RH-RhoGEF subfamily gets phosphorylated at Tyr-738 by Janus kinase 2 (JAK2) to positively regulate its GEF activity [57]. The presence of the phosphoserine residue suggests that K0513 may be required to undergo phosphorylation to perform its GEF function or other cellular processes.

*Escherichia coli* has been considered a pioneering host for recombinant protein production. Using *E. coli* remains a popular choice when attempting the production of a new protein because of the existing extensive knowledge on its genetics and physiology, and the number of genetic engineering tools adapted to *E. coli*. In this study, codon harmonization was applied to the human protein, K0513, through adjustment of the codon usage bias, adjustment of the GC content and removal of repeat sequences [58]. Codon harmonization by synonymous substitution of the coding sequence is designed to increase the expression level of the protein of interest [58]. K0513 was produced as a soluble recombinant protein. Its solubility may suggest cytosolic localisation (Fig 5) [6].

The fluorescence data provided preliminary insights on the tertiary structural features of K0513. Denaturing the protein with urea and guanidine-HCl resulted in a significant red shift of the intensity maxima in a concentration dependent manner (Fig 7). This reveals that the fluorescing Trp residues are located in a nonpolar hydrophobic interior [59]. This is because Trp has two nearby isoenergetic excitation states, ^1^L_a_ and ^1^L_b_ [60] which differs with sensitivity to solvent. In hydrophobic environments, ^1^L_b_ dominates the Trp emission and display a blue shift of Trp fluorescence due to an increase in non-polarity [61]. In contrast, denatured proteins with solvent exposed Trp residues displays a red shift [62]. The increase in the concentration of denaturants resulted in a significant red shift in the wavelength maxima (Fig 8), this shift suggests the exposure of Trp residues that are positioned within the hydrophobic interior [59]. This further suggests that, in its native state, K0513 exists as a globular protein. K0513 depicted no significant conformational changes when exposed to ATP, ADP, GTP nor GDP which suggests that K0513 may not be binding these nucleotides directly (Fig 7). This finding was supported by investigating the effect of nucleotides on the conformation of K0513 using limited proteolysis. The limited proteolysis assay showed that K0513 is a proteolytically susceptible molecule, however, its conformation remains unperturbed in the presence of nucleotides (Fig 9). K0513 was detected in a series of human cell lines, including lung carcinoma, fibrosarcoma and cervical adenocarcinoma cells, which could suggest its involvement in various pathways related to tumorigenesis, and possibly the importance of characterizing the function of K0513.

The *in silico* and biophysical analyses reported on in this study suggest that K0513 is involved in the Rab3a GTPase signalling pathway. Furthermore, due to lack of of significant conformational response to nucleotides, we suggest that K0513 may facilitate nucleotide exchange of Rab3a, by functioning as a GEF, inducing conformational changes in the Rab GTPase, and thereby activating the Rab3a. GEFs are potential targets for cancer therapy due to their role in many signalling pathways, especially in cell proliferation. For instance, several cancers result from mutations in the MAPK/ERK pathway that leads to proliferation. The GEF SOS1 activates Ras, whose target is the kinase Raf. Raf is a protooncogene due to the numerous mutations it exhibits in various cancers [63,64]. The Rho GTPase Vav1, which could be activated by the GEF receptor, has been shown to promote tumour proliferation in pancreatic cancer; GEFs represent possible therapeutic targets as they can potentially play a role in regulating these pathways through their activation of GTPases.

## 5. CONCLUSIONS

This is the first study to describe the expression, purification and characterisation of human K0513. Biophysical and *in silico* studies conducted on human K0513 here suggest that it is a soluble, globular protein that possesses SBF2 and dDENN domains. This suggests a role for this protein as nucleotide exchange factor for Rab3a. This implicates K0513 in neurotransmitter signalling pathways, possibly facilitating the role of Rab3a which is involved in the transport of synaptic vesicles in the brain [65,66], thus suggesting a crucial role for this protein in cancer development. Not only is the *K0513* upregulated in glioblastoma patients, but *K0513* has also recently been highlighted as one of four genes in a model to predict prognosis of pancreatic cancer [7]. The current study findings are important for further elucidation of the role of K0513, particularly in signalling pathways associated with tumorigenesis.

## 6. ACKNOWLEDGEMENTS

We thank the Hair and Skin Research Unit in the Department of Dermatology at the University of Cape Town Medical School for expertise and use of their cell culture facility.

## 8. DATA AVAILABILITY

All relevant data are within the manuscript and its Supporting Information files.

## 12. SUPPORTING INFORMATION

**S1 Table.**
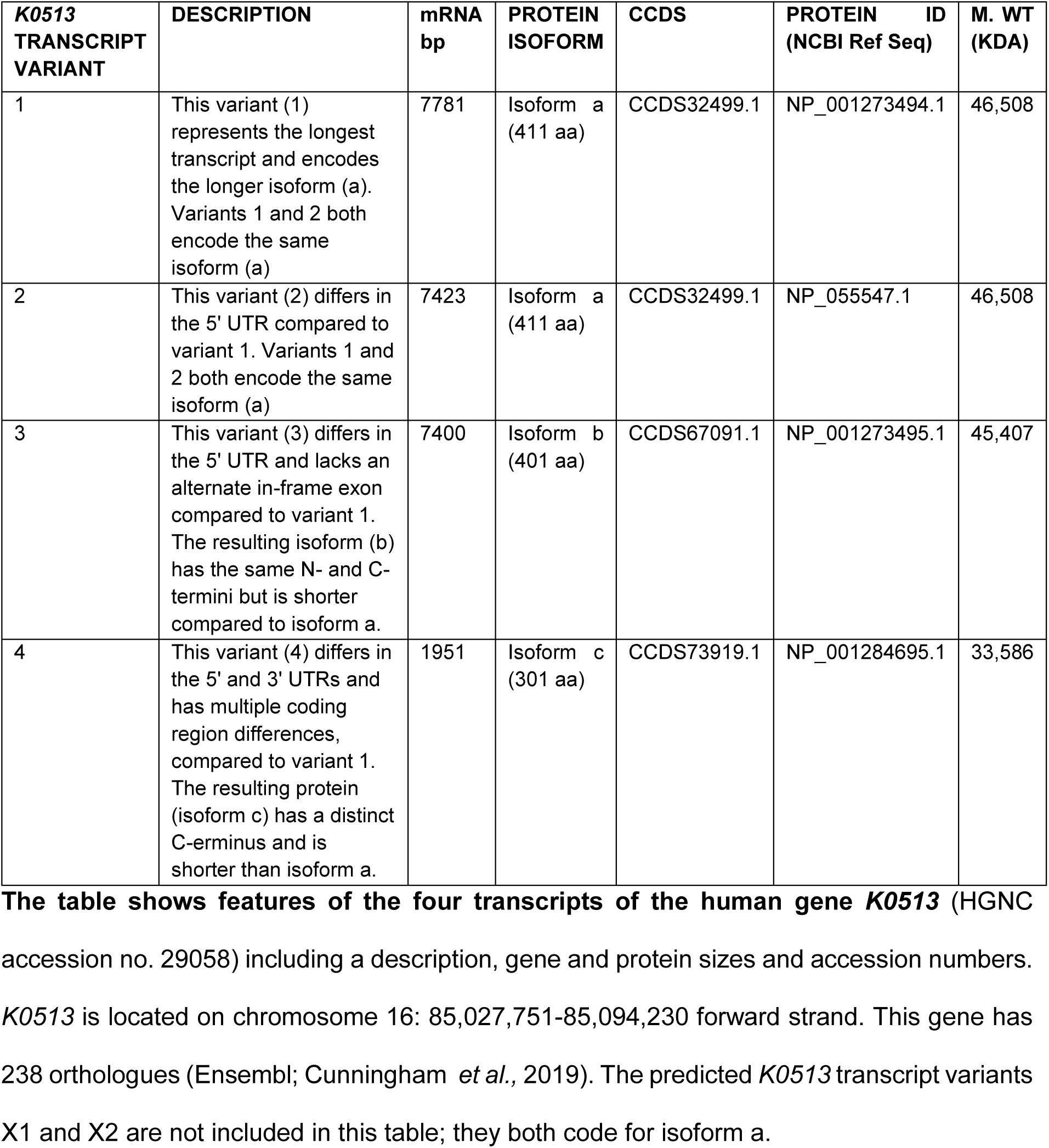
Summary of the seven transcript variants of human *K0513*. The table shows features of the four transcripts of the human gene *K0513* (HGNC accession no. 29058) including a description, gene and protein sizes and accession numbers. *K0513* is located on chromosome 16: 85,027,751-85,094,230 forward strand. This gene has 238 orthologues (Ensembl; Cunningham *et al*., 2019). The predicted *K0513* transcript variants X1 and X2 are not included in this table; they both code for isoform a.

| **K0513**<br>**TRANSCRIPT**<br>**VARIANT** | **DESCRIPTION** | **mRNA**<br>**bp** | **PROTEIN**<br>**ISOFORM** | **CCDS** | **PROTEIN ID**<br>**(NCBI Ref Seq)** | **M. WT**<br>**(KDA)** |
| --- | --- | --- | --- | --- | --- | --- |
| 1 | This variant (1) represents the longest transcript and encodes the longer isoform (a). Variants 1 and 2 both encode the same isoform (a) | 7781 | Isoform a (411 aa) | CCDS32499.1 | NP_001273494.1 | 46,508 |
| 2 | This variant (2) differs in the 5' UTR compared to variant 1. Variants 1 and 2 both encode the same isoform (a) | 7423 | Isoform a (411 aa) | CCDS32499.1 | NP_055547.1 | 46,508 |
| 3 | This variant (3) differs in the 5' UTR and lacks an alternate in-frame exon compared to variant 1. The resulting isoform (b) has the same N- and C-termini but is shorter compared to isoform a. | 7400 | Isoform b (401 aa) | CCDS67091.1 | NP_001273495.1 | 45,407 |
| 4 | This variant (4) differs in the 5' and 3' UTRs and has multiple coding region differences, compared to variant 1. The resulting protein (isoform c) has a distinct C-terminus and is shorter than isoform a. | 1951 | Isoform c (301 aa) | CCDS73919.1 | NP_001284695.1 | 33,586 |

**S2 Table.**
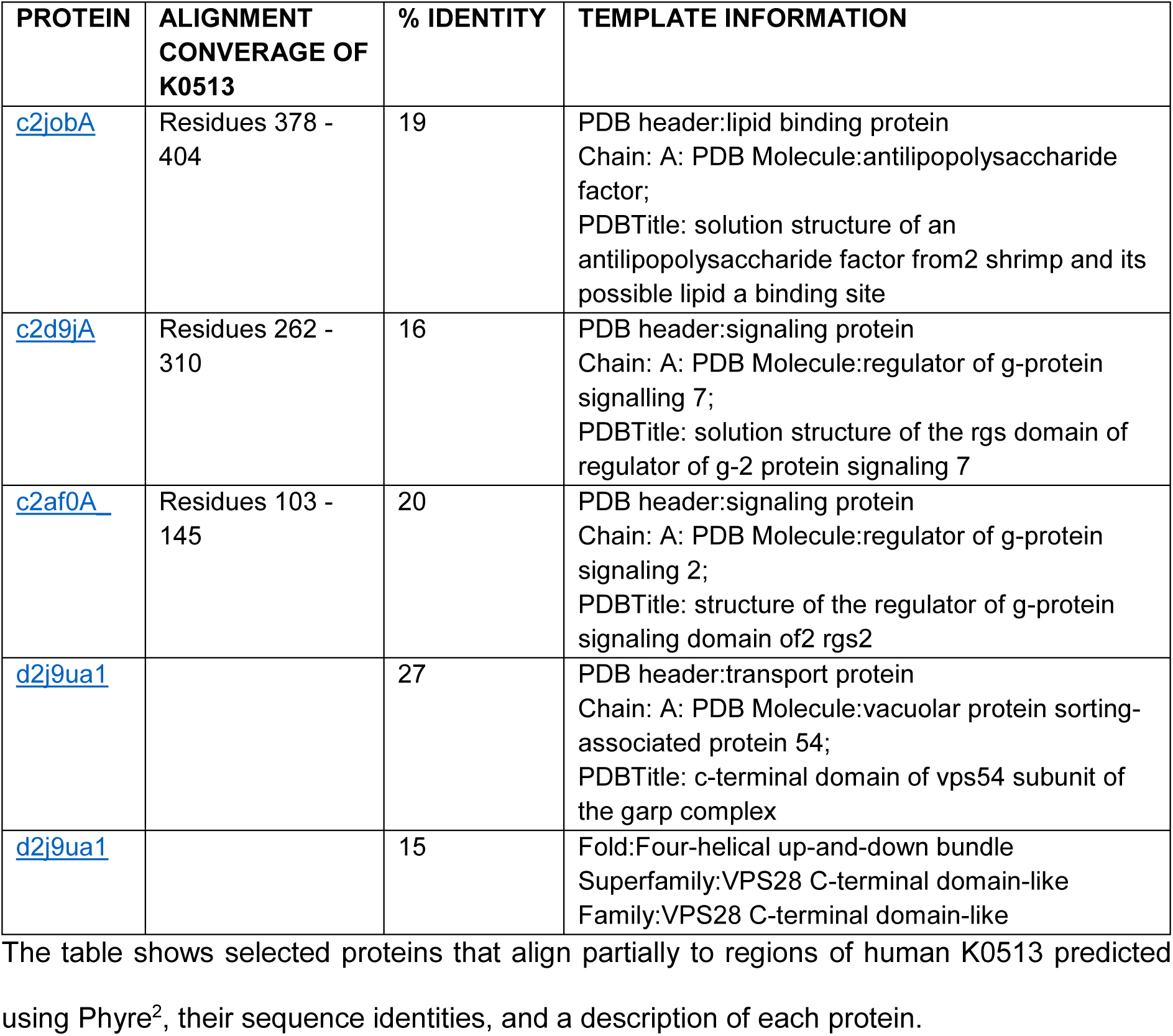
Proteins of interest that aligned to regions of K0513 as predicted by Phyre^2^. The table shows selected proteins that align partially to regions of human K0513 predicted using Phyre^2^, their sequence identities, and a description of each protein.

**S3 Table.**
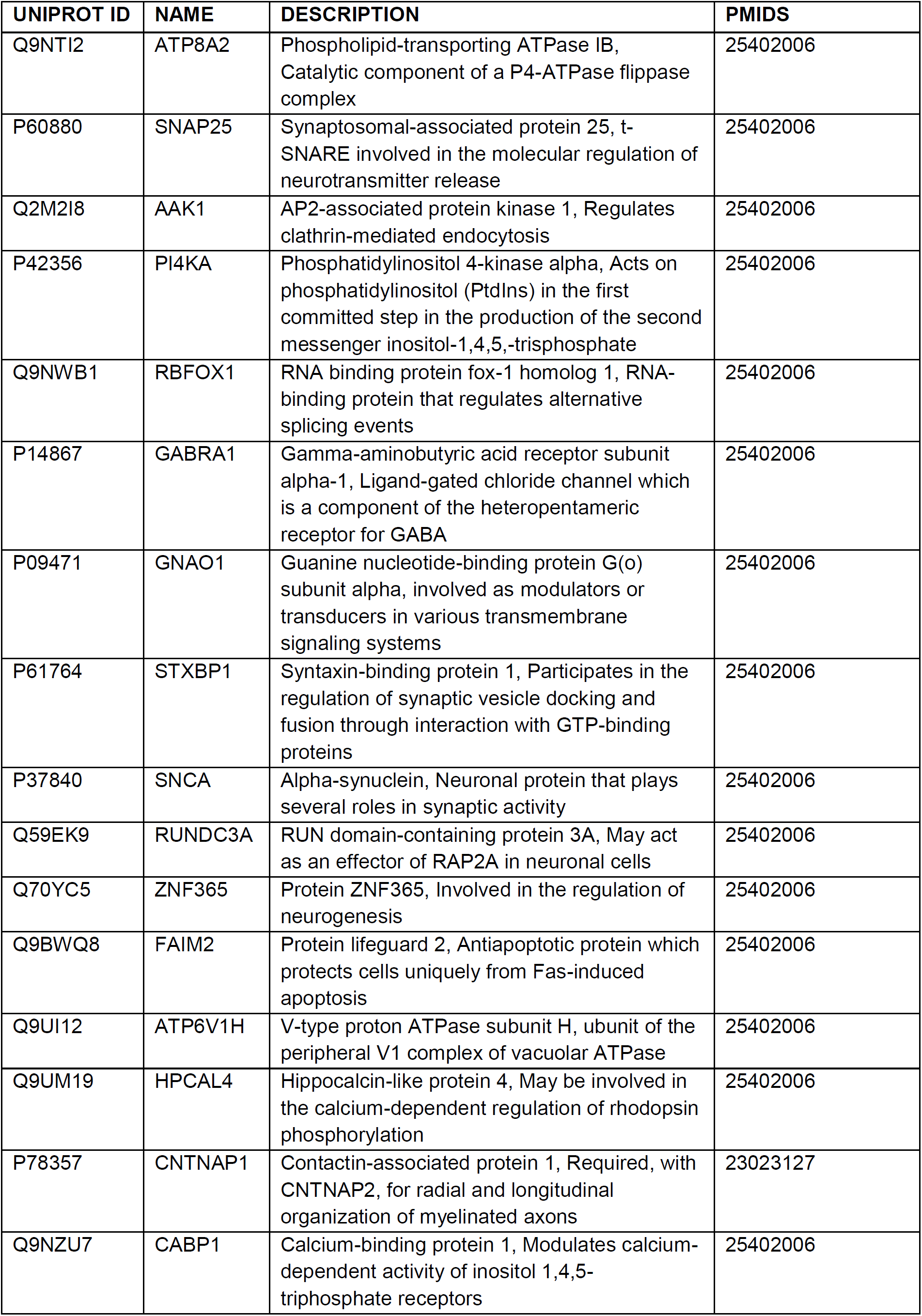
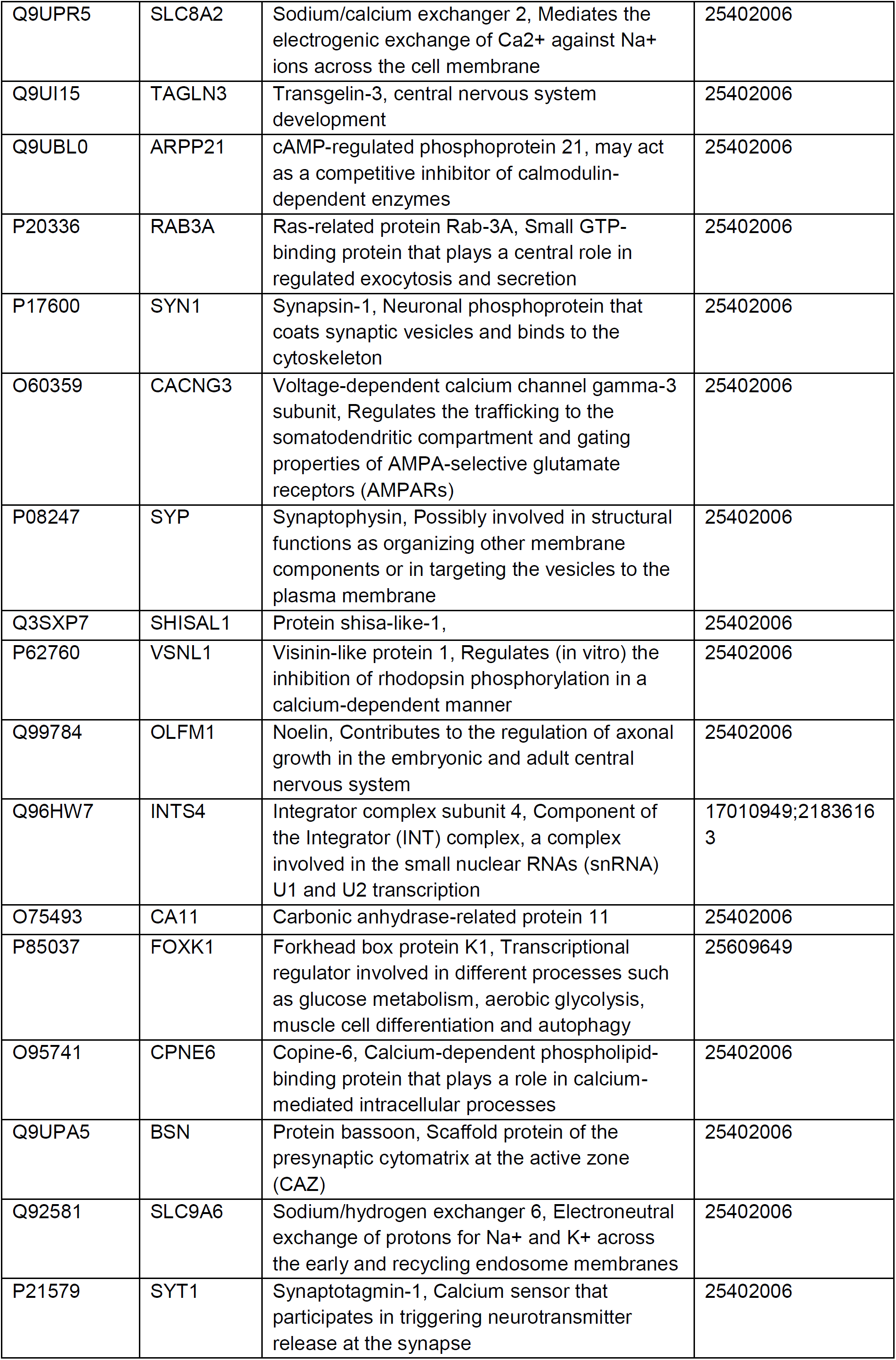
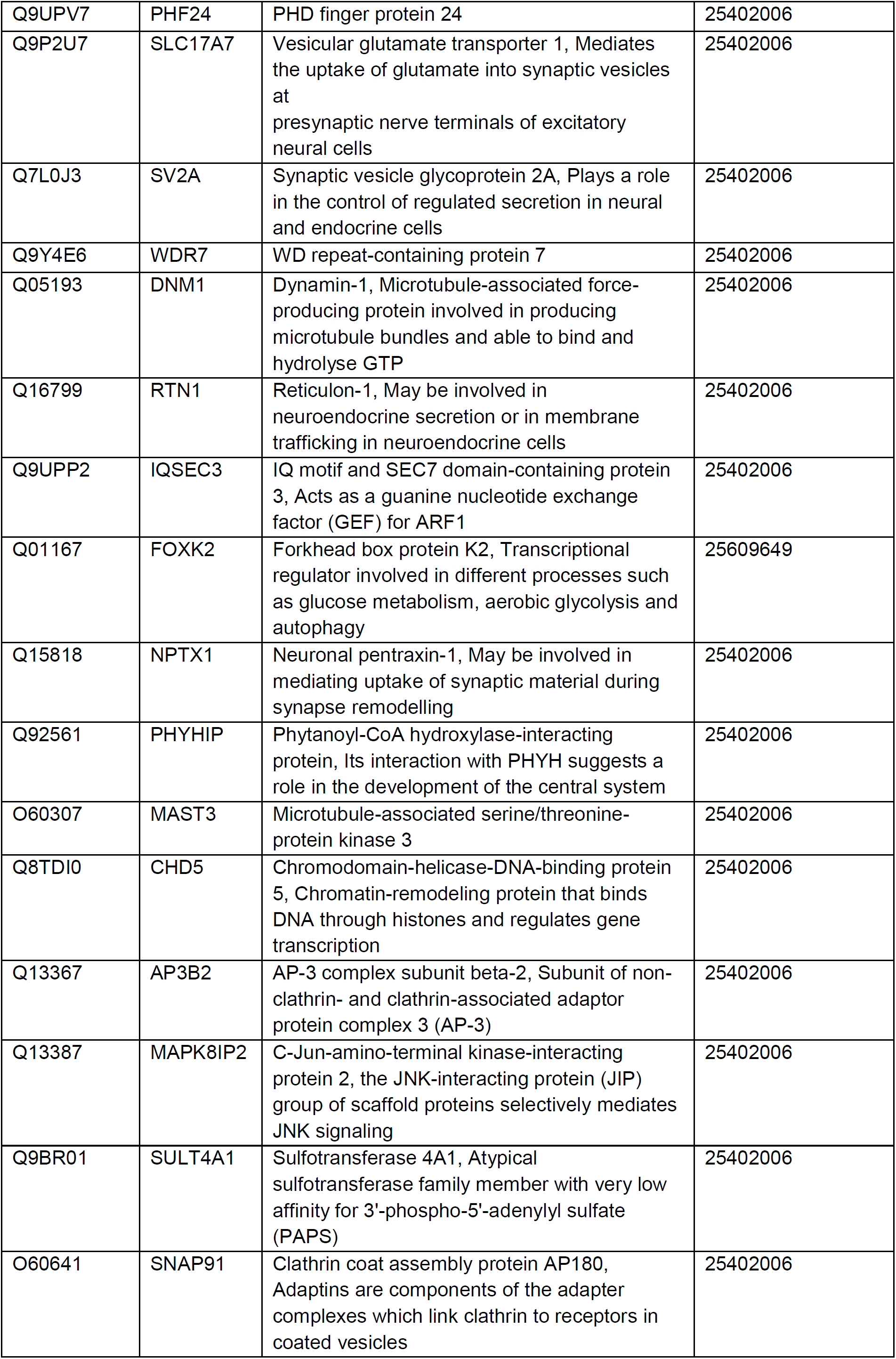
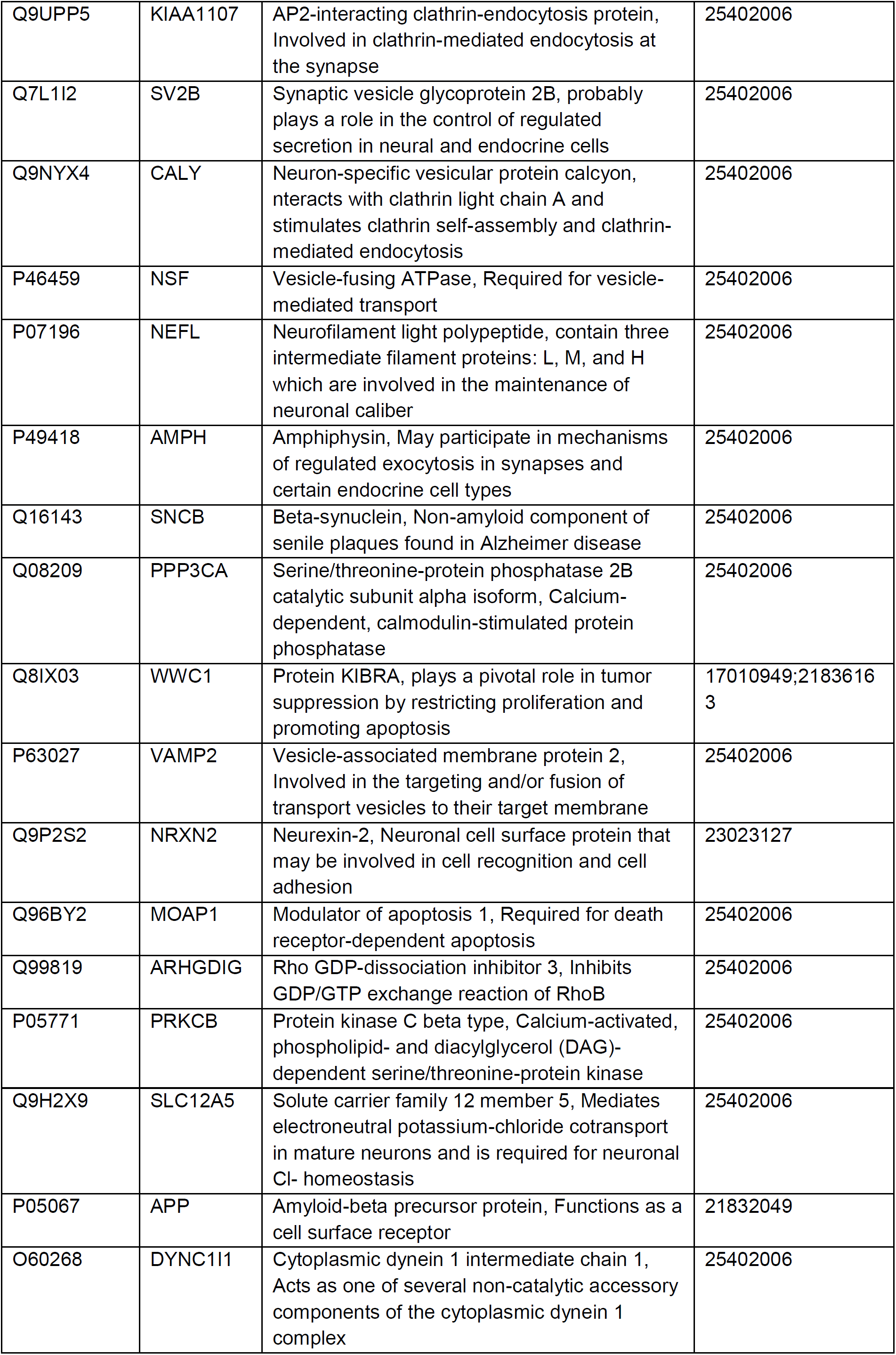
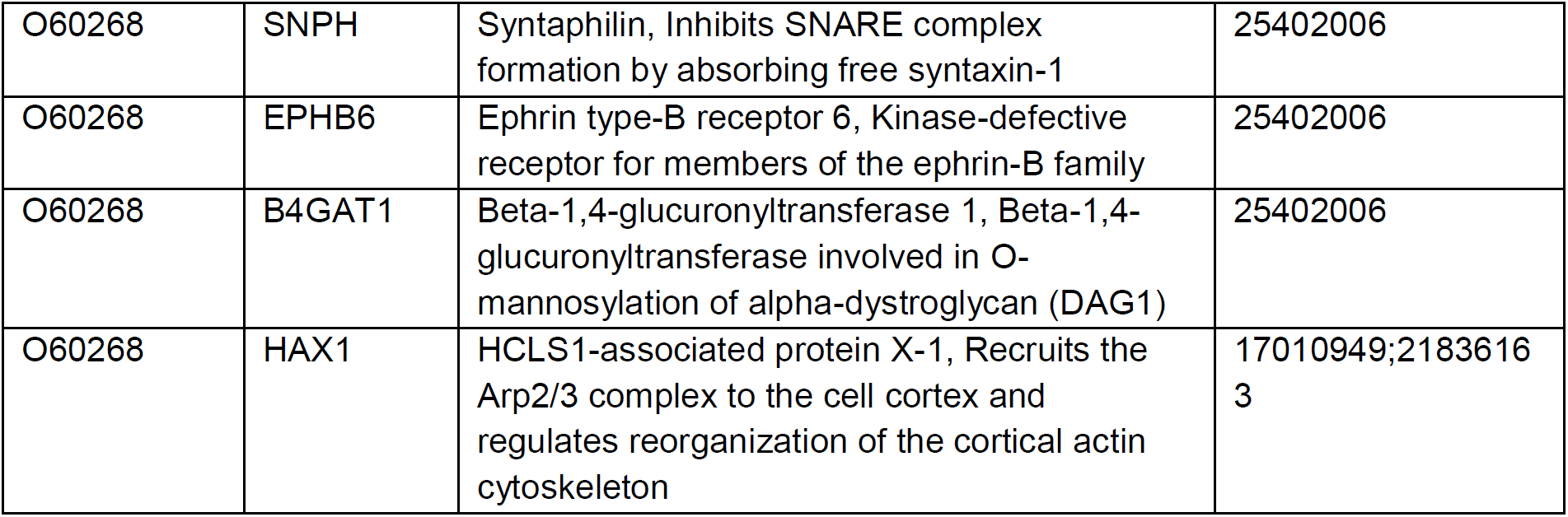
The predicted interactors of human K0513.

**S1 Fig.**
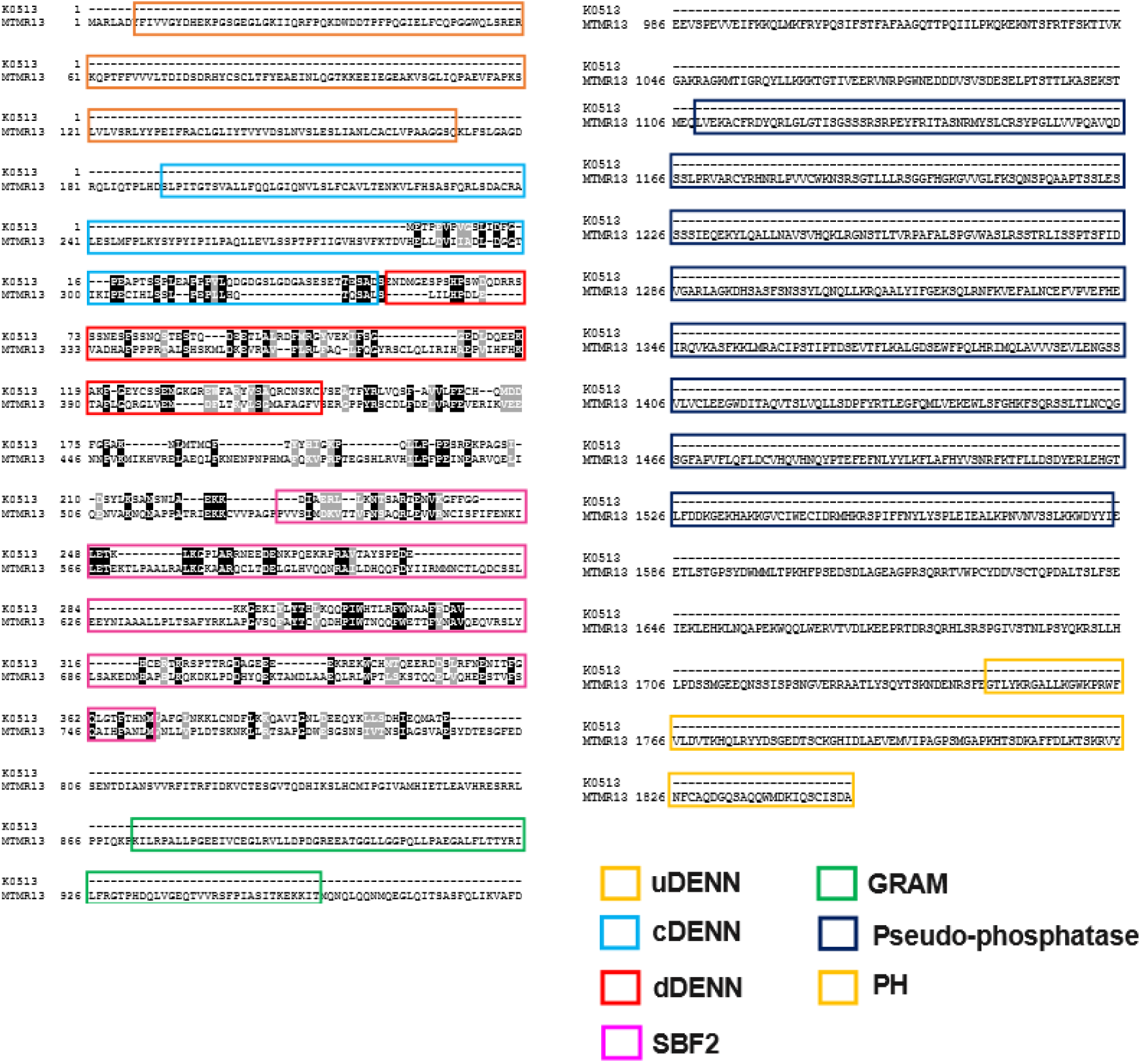
Pairwise sequence alignment of human K0513 and MTMR13. Pairwise sequence alignment of human K0513 (Accession: NP_055547.1) and human MTMR13 (Accession: Q86WG5.1). The uDENN (Olive green) is situated upstream of the DENN (green) followed by the dDENN (light green) domain. The hypothetical position of the SBF2 is represented by red, the GRAM (Glucosyltransferase/Rab-like GTPase activator/myotubularin domain) domain is represented by purple followed by the pseudo-phosphatase (Blue), and lastly the PH (Plenkin homolog) situated towards the C-terminal. The alignment was generated using EMBL-EBI Pairwise Alignemnt Tool (https://www.ebi.ac.uk/Tools/).

**S2 Fig.**
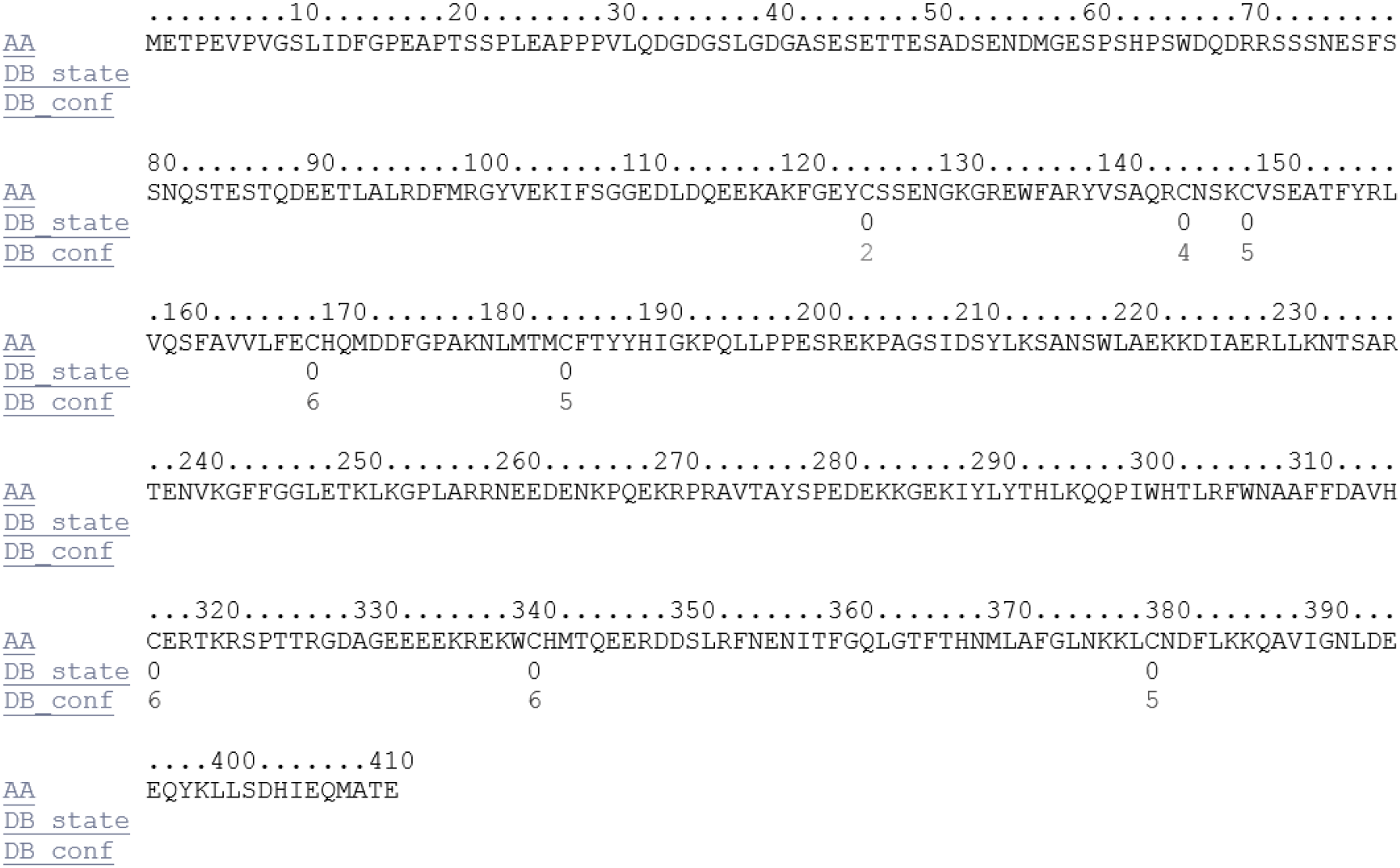
Prediction of human K0513 Cysteines disulfide bonding state and connectivity. No disulphide patterns were predicted for any of the eight cysteine residues present in human K0513 using the server DISULFIND. Symbol AA indicates amino acid sequence; DB_state indicates predicted disulfide bonding state where 1=disulfide bonded and 0=not disulfide bonded; and DB_conf indicates confidence of disulfide bonding state prediction where 0=low to 9=high.

